# Structural insight into outer membrane asymmetry maintenance of Gram-negative bacteria by the phospholipid transporter MlaFEDB

**DOI:** 10.1101/2020.06.04.133611

**Authors:** Xiaodi Tang, Shenghai Chang, Wen Qiao, Qinghua Luo, Yuejia Chen, Zhiying Jia, James Coleman, Ke Zhang, Ting Wang, Zhibo Zhang, Changbin Zhang, Xiaofeng Zhu, Xiawei Wei, Changjiang Dong, Xing Zhang, Haohao Dong

**Affiliations:** State Key Laboratory of Biotherapy and Cancer Center, National Clinical Research Center for Geriatrics, West China Hospital, Sichuan University and Collaborative Innovation Center of Biotherapy, Chengdu 610041, China; Department of Pathology of Sir Run Run Shaw Hospital, and Department of Biophysics, Zhejiang University, School of Medicine, Hangzhou, Zhejiang 310058, China; Center of Cryo Electron Microscopy, Zhejiang University, Hangzhou, Zhejiang 310058, China; College of Life Science, Sichuan University, Chengdu 610041, China; Biomedical Research Centre, Norwich Medical School, University of East Anglia, Norwich Research Park, Norwich, NR4 7TJ, UK

**Author notes:** These authors contributed equally to this work (Xiaodi Tang, Shenghai Chang, Wen Qiao, Qinghua Luo, Yuejia Chen). Correspondence (H.D.), (X.Z.), (C. D.).

## Abstract

The asymmetric phospholipid outer membrane (OM) of Gram-negative bacteria serves as the first line of defense against cytotoxic substances such as antibiotics. The Mla pathway is known to maintain the lipid asymmetry of the OM by transporting phospholipids between the inner and outer membranes. Six Mla proteins MlaFEDBCA are involved, with the ABC transporter MlaFEDB acts through a mechanism yet to be elucidated. Here we determine cryo-EM structures of MlaFEDB in apo, phospholipid-, ADP- or AMP-PNP-bound state to 3.3-3.75 Å resolution and establish a proteoliposome-based transport system containing MlaFEDB, MlaC and MlaA/OmpF to reveal the transport direction of phospholipids. Mutagenetic in vitro transport assays and in vivo sensitivity assays reveal functional residues which recognize and transport phospholipids as well as regulate the activity and structural stability of the MlaFEDB complex. Our work provides molecular basis for understanding the mechanism of the Mla pathway which could be targeted for antimicrobial drug development.

---

Gram-negative bacteria consist of two lipid membranes, the inner membrane (IM) and the outer membrane (OM), separated by an aqueous periplasmic space containing a thin peptidoglycan layer ^1^. Unlike the symmetrical IM, the OM is an asymmetric bilayer with an inner leaflet composed of primarily phospholipids (PLs) and an outer leaflet rich in lipopolysaccharides (LPS)^2–5^. The asymmetrical property of the OM facilitates its function as a selective barrier which prevents the entrance of cytotoxic molecules including antibiotics thus resulting in drug resistance^6–8^. Pathways that are responsible for trafficking OM components to maintain its asymmetry are therefore crucial biological processes for bacterial survival in environmental stress, which may be targeted for antimicrobial drug development^9–12^. Progress has been made in understanding the pathways for transporting and assembling OM components including the studies of β-barrel assembly machine (Bam) and LPS transport (Lpt) pathways^13,14,23,15–22^. The reported high-resolution structures and *in vitro* transport assays of seven Lpt transport proteins (LptA-G) allowed advanced understanding of the mechanisms how LPS is transported and assembled onto the outer leaflet of the OM ^5, 8, 32–34, 24–31^. On the other hand, the molecular mechanism of how PLs are trafficked between the IM and the OM to maintain the OM asymmetry remains unclear.

The maintenance of lipid asymmetry (Mla) pathway is believed to be important for maintaining the asymmetry and integrity of the OM by trafficking PLs between the IM and the OM^35–41^. Mutants of the Mla pathway in *Escherichia coli (E. coli)* or other pathogenic species show OM permeability and integrity defects and increased susceptibility to antibiotics^36–42^. Six proteins are involved in the Mla pathway, including an ATP-binding cassette (ABC) transporter MlaFE associated with an auxiliary periplasmic membrane protein MlaD and an auxiliary cytoplasmic protein MlaB, a periplasmic protein MlaC and an OM protein MlaA associated with an osmoporin OmpF/C^36,43,44^ (**Figure 1A**). Evidence has been provided from different studies to debate PLs transport direction of the pathway^35,36,43,45–49^. The Mla system was originally identified in *E. coli* as retrieving surface-exposed PLs in stressed cells to maintain OM lipid asymmetry and integrity. Mutants of the Mla pathway showed OM defects with increased SDS-EDTA sensitivity and accumulation of PLs in the outer leaflet of the OM^36^. A later study from molecular aspect analyzes how MlaA-OmpF/C complex in the OM removes PLs from the outer leaflet of the OM, backing up the hypothesis of the retrograde PL transport of the Mla system^47^. More recent studies, however, implicate a possible anterograde PL transport as observed accumulation of PLs in the IM upon destructing the Mla homologous pathway in *Acinetobacter baumannii*^48^. Another anterograde supporting study shows that the soluble periplasmic domain of MlaD passes PL to MlaC (anterograde direction) but not the other way around^49^. In either case, MlaC is believed to act as a chaperon in the periplasm to deliver PLs between MlaA and MlaD^42,45,49^. These studies may have provided clues for understanding the transport direction of the Mla pathway but also suggest a phenomenon that PLs are trafficked between MlaA and MlaD via MlaC without energy consumption, neglecting the role of the ABC transporter MlaFEDB in the pathway. MlaF, MlaE, MlaD and MlaB form a stable complex in the IM with a molecular ratio 2:2:6:2 ^45^. Two recent studies determined MlaFEDB cryo-Electron Microscopy (cryo-EM) structures of *E. coli*^45^ and *A. baumannii*^48^ at resolutions of 12 and 8.7 Å respectively; however these low-resolution structures did not reveal molecular details of the complex thus the role of MlaFEDB in Mla pathway remains unclear.

**Figure 1.**
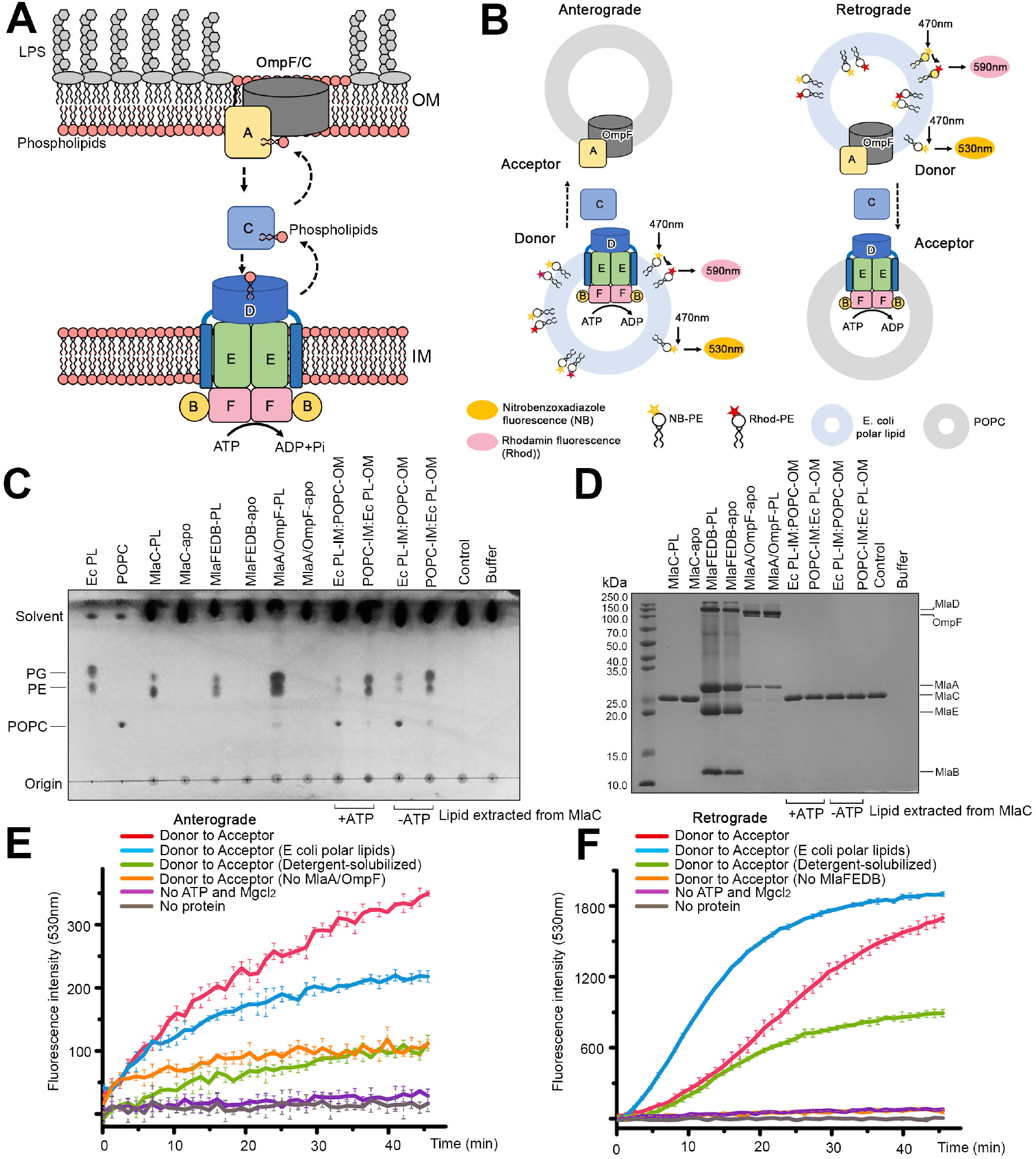
PL transport between the OM and IM proteoliposomes by the Mla pathway. (A) A scheme of the Mla pathway showing transport of PL between the OM and the IM involving the IM complex MlaFEDB, periplasmic protein MlaC and OM complex MlaA/OmpF. (B) Schematic diagram of the FRET based PL transport assay showing the anterograde (left) and retrograde (right) transport of PL. MlaFEDB or MlaA/OmpF complex was reconstituted into the donor or acceptor liposomes. The donor liposomes were constructed using E. coli polar lipid extract, nitrobenzoxadiazole (NB) labelled PE and rhodamine (Rhod) labelled PE. The acceptor liposomes were constructed using unlabeled 1-palmitoyl-2-oleoyl-glycero-3-phosphocholine (POPC). NB emits at 530nm (shown as orange) and Rhod emits at 590nm (shown as pink). (C) PL transport assay using unlabelled IM and OM proteoliposomes and MlaC in the presence of absence of ATP and MgCl_2_. All proteins used in the assay were in apo states. PL transported to MlaC were detected by TLC for retrograde and anterograde transport. PLs transported from OM proteoliposomes were more potent than from IM proteoliposomes. (D) SDS-PAGE of the proteins used for the assay in Figure C. (E and F) PL transport assay using fluorescent PE mixed donor and POPC acceptor proteoliposomes for FRET measurement of the NB signal at 530nm for anterograde (E) and retrograde (F) directions in the presence of 1mM ATP and 2mM MgCl_2_. The retrograde transport shows a higher FRET signal than the anterograde and is dependent on the composition of the IM proteoliposomes. Representative data of n≥3 experiments.

In this study, we used biochemical transport assays and cryo-EM structures to determine the transport direction and structural characteristics of MlaFEDB. Our transport assays reveal a predominant retrograde PL transport between the constructed IM and OM proteoliposomes; the high-resolution structures and the mutagenic in vivo and in vitro assays reveal functional residues for PL recognition and transportation as well as structural regulations of each subunit to the ATPase and transport activity of the MlaFEDB complex. Our findings provide insight into the mechanism of how the asymmetry of the outer phospholipid membrane is achieved and maintained in Gram-negative bacteria to resist environmental stress, which could be therapeutically targeted for antimicrobial drug development.

## Results

### MlaFEDB transports Phospholipids bidirectionally in vitro

We expressed MlaFEDCB operon from E. coli strain K-12 and obtained a membrane complex of MlaFEDB without MlaC. Purified MlaFEDB showed ATPase activity that was higher in *E. coli* polar lipids liposomes than in detergent micelles of n-dodecyl-beta-D-maltoside (DDM) and inhibitable by ADP or AMP-PNP (**Figures S1A-S1D**). To determine how Mla pathway transfers PL, we established an *in vitro* PL transport system using both purified MlaFEDB and MlaA/OmpF constructed IM and OM proteoliposomes respectively and systematically monitor the PL transport direction in the presence of MlaC (**Figure 1B**). To track the origin of transported PLs we differentiated the IM and OM proteoliposomes by using *E. coli* polar lipids containing phosphatidylethanalomine (PE), phosphatidylglycerol (PG), and cardiolipin (CL) or non-bacterial phospholipid 1-palmitoyl-2-oleoyl-glycero-3-phosphocholine (POPC). The reconstitution ratio and orientation for both MlaFEDB and MlaA/OmpF were determined to ensure IM and OM proteoliposomes were equally constructed (**Figures S1E and S1F**). We first determined PL transport to apo-MlaC by using thin layer chromatography (TLC) and detected major PE/PG or POPC transported from OM proteoliposomes, and minor PE/PG transported from IM proteoliposomes, which were both independent of ATP and MgCl_2_ (**Figures 1C and 1D**). It is consistent to the previous reports that MlaFEDB^49,50^ or MlaA/OmpF^47^ passively transport PLs to apo-MlaC *in vitro* as MlaC shows higher PL affinity. Interestingly, our apo-MlaFEDB and apo-MlaA/OmpF constituted proteoliposomes showed PL transport same as the PL-bound form (**Figures 1C, 1D, S1G and S1H**), suggesting that MlaFEDB and MlaA/OmpF not only pass PL to MlaC but also extract PLs from membranes. The retrogradely transported PL from the OM proteoliposomes was abolished by deleting MlaA Asn41 and Phe42 (MlaΔAsn41-Phe42) (**Figures S1G-J**) which was previously reported to be functional residues for retrograde transport of PLs^37^. The anterograde PL transport was not affected by the mutant MlaAΔAsn41-Phe42/OmpF (**Figures S1G and S1H**).

For quantification, we used fluorescent nitrobenzoxadiazole (NB) and rhodamine (Rhod) labeled PE (NB-PE and Rhod-PE) mixed with *E. coli* polar lipids for donor proteoliposomes and unlabeled POPC for acceptor proteoliposomes to systematically record the transport of fluorescent PE for a specific direction (anterograde=IM donor/OM acceptor; retrograde=IM acceptor/OM donor) (**Figure 1B**). Proteins used in FRET assays were not treated for apo-state unless stated. The results show an increasing NB signal over time for both anterograde and retrograde transport as a result that the labeled PE was transported out of the Fluorescence Resonance Energy Transfer (FRET) range between NB and Rhod. The retrograde transport is much more efficient than anterograde at the physiological ATP concentration of 1mM within the tested time course, which is consistent with the TLC result that the retrograde is predominant (**Figures 1E and 1F**). The NB or Rhod labeling on the head group of PE does not seem to have adverse effect on the transport as the fluorescent PE signal captured by MlaC showed similar trend comparing with the TLC results using unlabelled PLs (**Figure S1K**). Interestingly, in contrast to the TLC result, FRET recorded PL transport for both directions were ATP and MgCl_2_ dependent and inhibitable by ATP or AMP-PNP (**Figures S1L and S1M**). This suggests that the retrograde PL transport also involve the ATPase activity of MlaFEDB. Indeed, the rate of retrograde transport was hugely influenced by the composition of the IM proteoliposomes (**Figure 1F**). Transport assay using empty IM proteoliposomes containing no MlaFEDB almost abolished retrograde transport FRET signal whereas changing IM proteoliposomes from POPC to *E. coli* polar lipids massively accelerated retrograde transport (**Figure 1F)**.

For retrograde transport, MlaFEDB is expected to import PLs from MlaC, which returns the bioavailability of apo-MlaC for further retrograde transport. FRET transport assay using detergent-solubilized MlaFEDB as acceptor showed reduced retrograde transport at later time comparing with that of using proteoliposome acceptor, suggesting that the import action of MlaFEDB is required for efficient and continuous retrograde transport of the Mla pathway (**Figure 1F**). On the other hand, TLC results show that anterograde PL transport does not require the ATPase activity of MlaFEDB thus the ATP-dependent anterograde FRET signal was probably mainly contributed by the reflux of unlabeled PLs from OM proteoliposomes (**Figure 1C**). Anterograde transport using empty OM proteoliposomes or detergent MlaA/OmpF as acceptor show similar reduced FRET signal however not as low as the basal level (**Figure 1E**). Although weak, FRET scan revealed ATP-independent anterograde signal in the presence of apo-MlaC (**Figure S1N**) or when blocking the retrograde transport by using the mutant MlaAΔAsn41-Phe42/OmpF (**Figures S1I and S1J**). This suggests that anterograde transport requires un-clogged transport path for PLs to diffuse through and the retrograde transport of PL is prioritized by MlaFEDB. To test whether PL was further transported from MlaC to the OM for anterograde we scanned fluorescent signals in isolated MlaA/OmpF proteoliposomes for any transported NB-PE or Rhod-PE from IM proteoliposomes. The result shows both increased NB signal (540nm) and Rhod signal (590nm) in the OM proteoliposomes (**Figure S1O**). However, we also detected large amount of MlaC co-purified with the MlaA/OmpF proteoliposomes even after several washes with buffer (**Figure S1P**). Although no direct evidence to support that PLs were transported into the OM, the results at lease show the tendency of MlaC carrying anterogradely transported PLs to associate with the OM. To summarize, our *in vitro* transport assays suggest an ATP-dependent retrograde transport of PL from MlaA/OmpF proteoliposomes to MlaFEDB proteoliposomes and a passive anterograde transport of PLs from MlaFEDB proteoliposomes to MlaC. Although no evidence was shown for anterograde PL transport between MlaC and MlaA/OmpF, our findings implicate that MlaFEDB is a bidirectional transporter and it functions predominantly for the retrograde PL transport. The transport direction of MlaFEDB may be determined by the concentration of PLs carried in the MlaC pool, charged by the upstream of the retrograde transport but it requires further investigation to confirm.

### The architecture of four MlaFEDB cryo-EM structures

To understand the transport mechanism of MlaFEDB, we investigated the molecular structures of the complex. Purified MlaFEDB protein was incubated with *E. coli* polar lipid extract (PLs), AMP-PNP or ADP for 1hr at room temperature to obtain PL-bound, AMP-PNP-bound and ADP-bound states, respectively. Cryo-EM structures of MlaFEDB in apo, PL-bound, AMP-PNP-bound and ADP-bound states were determined to 3.4, 3.3, 3.4 and 3.75 Å resolutions, respectively (**Figures S2-S5, Table S1**). The high-resolution maps allowed us to build models with confidence and assign amino acids precisely based on clear densities of their side chains (**Figures S2-S5, Methods**). The architecture of MlaFEDB resembles the shape of a “oil lantern” with a unique “lampshade” like structure of hexameric MlaD that form a hexameric disc covering from the periplasmic side and six transmembrane helices surround the transmembrane core of the dimeric MlaE (**Figures 2A and 2B**). The dimeric MlaF in the cytoplasm constitutes the base of the complex with an auxiliary protein MlaB associated with each MlaF subunit (**Figures 2A-E**). Each MlaE unit has five vertical transmembrane (TM) segments (TM1-5) and one N-terminal cytoplasmic elbow helix that is parallel to the IM plane, showing a novel topology (**Figure 2F**). Between TM segments, each MlaE forms two periplasmic loops (loop 1 and loop 2) and two cytoplasmic turns (turn #1 and turn #2). The periplasmic loop 2 between the TM3 and TM4 carries an a-helix that makes interactions with the periplasmic domains of MlaD (**Figures S6A and S6B**). The cytoplasmic turn #1 between the TM2 and TM3 contains a coupling helix that interacts with the dimeric MlaF (**Figures S6A and S6C**). Two MlaE units form a dimer of transmembrane domains (TMDs) through the interactions between the respective TM1 and TM5 of the two protomers (**Figures 3A-C**). Each MlaE is evenly surrounded by three TMs that come from three adjacent MlaD subunits of the hexamer, making interactions with the elbow helix, TM1 and TM3 of MlaE (**Figures 2B, 3C, S6D, and S6E**). On the cytoplasmic side, the dimeric MlaF form the nucleotide-binding domains (NBDs), each of which contains a helical and a RecA-like domain sharing folds with other reported ABC transporter ^51,52^ (**Figure S6A**). Two MlaB units interact with MlaF on the far ends through their respective a-helices (**Figures 2A-E**). The nucleotide (AMP-PNP or ADP) and phospholipid-bound structures of MlaFEDB show similar conformations to the apo structure with a rootmean-square deviation (RMSD) of 1.0 Å over 2021 aligned residues, 1.43 Å over 1873 aligned residues, and 1.25 Å over 1928 aligned residues respectively (**Figures 7A-D**). Nevertheless, these structures reveal distinct substrate and nucleotide binding details for understanding the transport mechanism of MlaFEDB.

**Figure 2.**
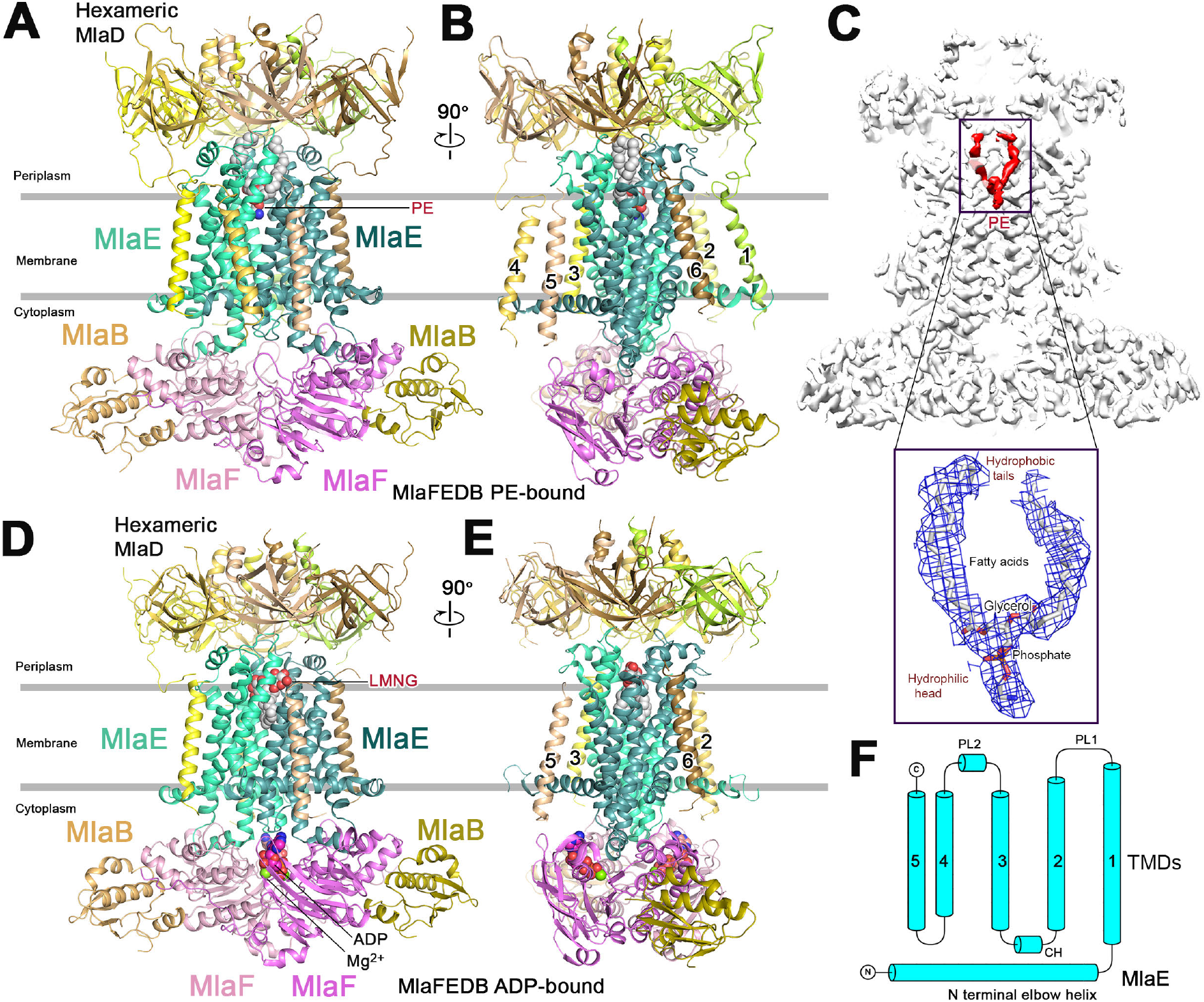
Architectures of PE- or ADP-bound MlaFEDB complexes. (A) Cartoon representation of the PE-bound MlaFEDB structure. The hexameric MlaD are in yellow-brownish colours. Two MlaE molecules are in green and dark green. Two MlaF molecules are in violet and light pink. MlaB molecules are in olive and light orange. PE is shown in sphere. (B) PE-bound MlaFEDB complex rotated 90° along y-axis relative to the left panel. The TM domains of MlaD are numbered. (C) Cryo-EM map of PE-bound MlaFEDB structure. PE is in red. The density map of the bound PE at the lower panel. (D) Cartoon representation of ADP-bound MlaFEDB complex. The colour scheme is the same as A. LMNG and ADP are shown in sphere. (E) Rotation of 90° along y-axis relative to the left panel. Only four TM domains of MlaD are clearly visible and numbered. (F) Topology of MlaE, containing an elbow a-helix and five transmembrane segments.

**Figure 3.**
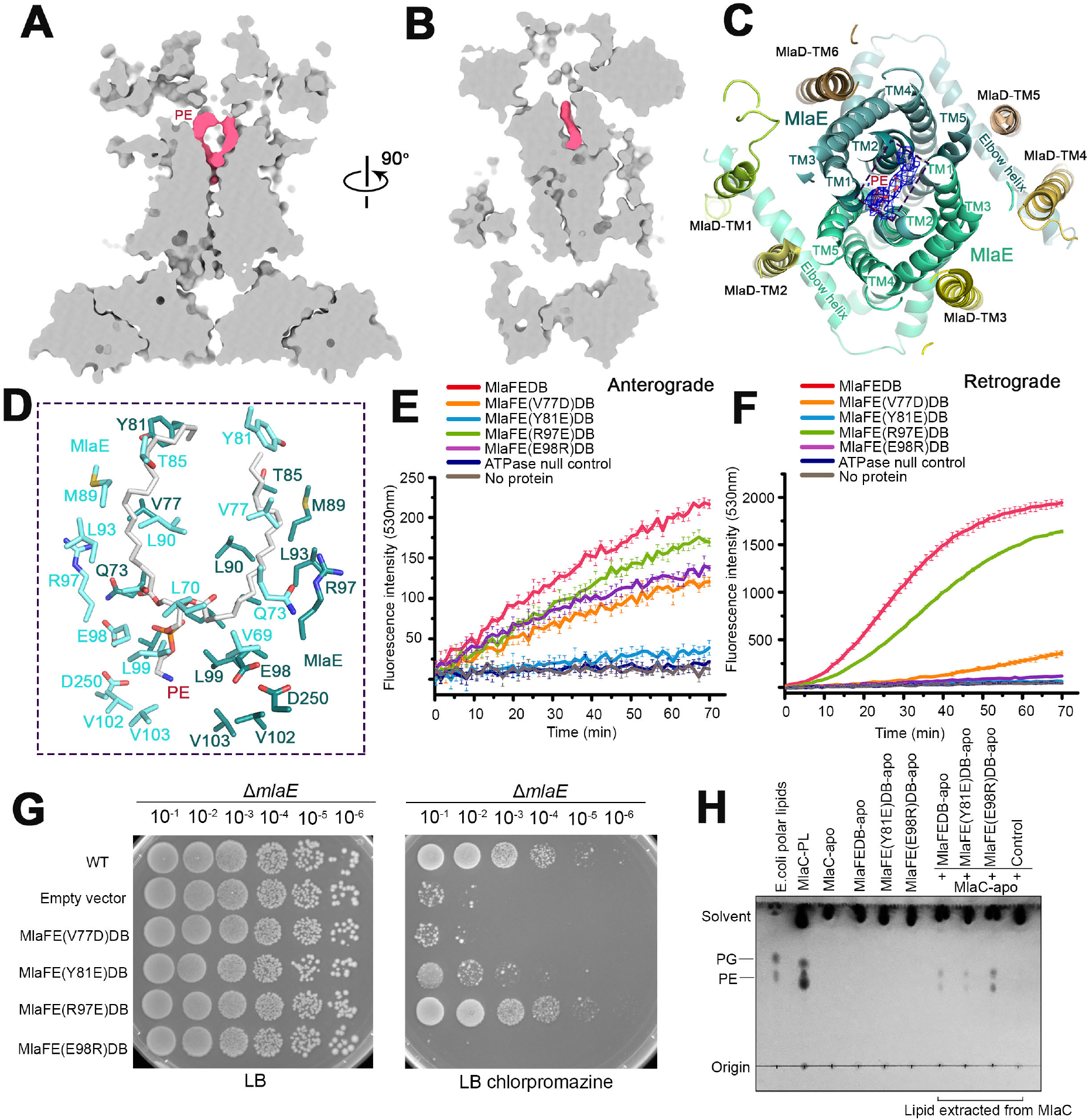
MlaFEDB recognition of PE. (A) Cross-sectional view of the cryo-EM map of PL-bound MlaFEDB. The density for PL is in pink in the cavity of the transporter. (B) Cross-sectional view of the cryo-EM map of PL-bound MlaFEDB rotated 90° along y axis relative to A. (C) Slab view of the TM segments of the MlaFEDB from the top. The dimeric MlaE form the PE binding cavity, and the six TM segments of MlaD surrounded MlaE. The colour scheme of MlaFEDB is the same as Figure 2A. (D) Cavity residues interacting with the bound PE. (E and F) FRET assays measuring PL transport for anterograde (E) and retrograde (F) directions in response to mutation of PE binding residues on MlaE. Single MlaE mutant Y81E abolishes anterograde transport while Y81E and E98R abolish the retrograde PL transport. Representative data of n≥3 experiments. (G) Cell based assays of MlaE’ s PL bound residue mutants using the MlaE knockout strain. The assays were carried on LB plate without (left) or with 120 μg ml^-1^ chlorpromazine (right). (H) Anterograde PL transport assay using apo-MlaFE(Y81E)DB or apo-MlaFE(E98R)DB constituted IM proteoliposomes and apo-MlaC in the absence of ATP. PL transported to MlaC was resolved by TLC. MlaFE(Y81E)DB reduced while MlaFE(E98R)DB increased the anterograde transport comparing to MlaFEDB.

### MlaFEDB retrogradely transports PL in a head-down orientation

MlaFEDB contains a V-shaped TM cavity created by the TM1, 2 and 5 of the MlaE dimer which expands into the periplasm and creates a continuous pathway with the central channel of the periplasmic hexameric disc of MlaD (**Figures 2A-C**). Hydrophobic residues of MlaE face inside and constitute most of the cavity and only a few numbers of polar and charged residues decorated at the midway of the cavity. The cavity is formed by MlaE residues Tyr81, Met89, Leu78, Val77, Leu90, Arg97, Glu98, Ser94, Gln73, Leu70, Ile66, Leu99, Val102, Val103 and Leu247 (**Figure 3D**). In the MlaFEDB PL-bound structure, a density for a L-a-phosphatidylethandamine (PE) is observed in the cavity (**Figures 2C, 3A, 3B, and S3F**). The polar phosphate head of the bound PE is facing downward surrounded by the residues at the midway of the cavity including Glu98, Leu70, Leu99 while the two acyl chains extend towards the periplasm interacting with the hydrophobic residues in the upper cavity including Gln73, Val77, Met89, Tyr81, Leu93, Arg97 (**Figure 3D**). To test whether these substrate binding residues are important for PL transport, we made single mutants Tyr81Glu, Val77Glu, Arg97Glu and Glu98Arg and performed *in vitro* PL transport assay and *in vivo* cellular sensitivity assay (**Figures 3E-3H**). Compound chlorpromazine as a membrane stress or has been used in reports for cellular sensitivity assays ^47^. The results showed that the mutations Tyr81Glu and Glu98Arg abolished the retrograde PL transport and Glu98Arg also severely reduced cellular survivability to chlorpromazine (**Figures 3F, 3G, and S8H**) suggesting that these residues are involved in retrograde PL transport. For anterograde FRET, the transport is affected mainly by Tyr81Glu but not Glu98Arg (**Figure 3E**). TLC transport assay using MlaFEDB proteoliposomes and apo-MlaC in the absence of ATP shows decreased anterograde PL transport by the mutant MlaFE(Tyr81Glu)DB but increased transport by the mutant MlaFE(Glu98Arg)DB (**Figures 3H, S8I,a nd S8J**). Glu98 is present at midway of the cavity interacts with the polar head of the “head-down” orientated PE (**Figure 3D**), suggesting that the head-down orientated PL in the cavity is being transported retrogradely and the way that MlaFEDB transports PL for anterograde may be different. In the apo MlaFEDB structure, some extra densities are observed in the cavity, which are identified to be two Lauryl Maltose Neopentyl Glycol (LMNG) molecules (**Figures S8A-G**). These two detergent molecules positioned one on top of the other, with their sugar heads pointing toward the periplasm and the two hydrophobic tails pointing towards the bottom of cavity (**Figure S8**). The top LMNG molecule locates at the periplasmic interface between MlaE and MlaD with its hydrophobic tails interacting with the hydrophobic residues in the upper cavity including Tyr81, Met89, Leu78, Val77 and Leu90, while the lower LMNG molecule has its acyl tails interacting with the hydrophobic residues in the lower cavity including Leu70, Ile66, Leu99 and Val103 (**Figure S8G**). In the ADP-bound structure, there is also one LMNG molecule bound at the lower cavity, which orientates and interacts similarly as the one in the apo MlaFEDB structure (**Figures 2D, 2E, S4F, and S9A**). The orientation of the bound detergent molecules is inverted from that of the bound PE molecule, raising a possibility that PL could also be accommodated in the cavity of MlaFEDB in a head-up orientation. It is worth noting that both ADP and AMP-PNP bound structures contain no ligand binding in the upper cavity while the non-nucleotide bound structures contain PL or detergent molecules, suggesting that nucleotide binding to MlaFEDB may induce an exit of the bound ligands.

### MlaD regulates MlaFEB ATPase and transport activity

The C-terminal periplasmic domain of MlaD form an SDS-resistant hexameric disc interacting with the TMDs of the complex through the periplasmic loop 1 and 2 of MlaE at the interface (**Figures S6A and S6B**). Each MlaD contains a TM segment at the N-terminus. Three TM segments from adjacent MlaD subunits interact with each MlaE protomer at the conserved Gly10 and Gly21 of the elbow helix and the surface residues of TM1 and TM3 (**Figures S6D and S6E**). In the apo and PL-bound MlaFEDB structures six TM segments from MlaD are observed (**Figures 2A and 2B**) while in the ADP and AMP-PNP-bound structures only clear densities for four TM segments were shown (**Figures 2D, 2E, S9B, and S9C**). The one that interacts with the elbow helix was not resolved (ADP-bound structure) or partially resolved (AMP-PNP-bound structure) (**Figures 2D, 2E, S9B, and S9C**) suggesting that the TM segments of MlaD are conformational flexible and may be involved in transport action of MlaFEDB. To test this, we deleted the gene from Gln2-Asn30 encoding for the TM helix of MlaD. Truncated MlaD(ΔTM) was no longer purified in complex with MlaFEB suggesting that the TM segments are required for stable complex formation (**Figure S10F**). Despite that, isolated MlaD(ΔTM) or full length MlaD forms hexamer regardless the presence or absence of the TM segments, and the resulted full length MlaD hexamer could form stable complex with MlaFEB *in vitro* (**Figure S10A**). Compared with MlaFEDB, MlaFEB constructed IM proteoliposomes showed little ATPase activity (**Figure S10B**) and completely abolished PL transport in both directions, suggesting that the presence of MlaD is essential for the transport activity of MlaFEB for both directions (**Figures S10C and S10D**). Interestingly, in detergent micelles MlaFEB shows higher ATPase activity than MlaFEDB (**Figure S10B**), implying the regulatory effect of MlaD to the ATPase activity of MlaFEB. Introducing soluble MlaD(ΔTM) to either detergent or liposome constituted MlaFEB showed no recovery of PL transport or ATPase activity whereas *in vitro* incubated full length MlaD was able to rescue both activities (**Figures S10B-S10D**). These results suggest that the effect of MlaD on the activity of MlaFEDB is only realized in the presence of the TM segments which bring MlaD and MlaFEB into a stable complex, revealing the importance of the TM segments of MlaD.

In the periplasmic domain, residues Leu143-Gly153 of each MlaD form a short helix, which associates together to create a channel in the center of the MlaD hexamer (**Figures 4A and 4B**). Truncated MlaD(ΔL143-G153) lost its hexameric form (**Figure S10F**). Hydrophobic residues Leu143, Leu146, Ile147 and Phe150 face inside the channel and the residue Tyr152 is on the top of the channel (**Figure 4B**). Single MlaD mutants Leu143Glu, Ile147Glu or Tyr152Glu severely impaired PL transport in both directions as well as cellular survivability to chlorpromazine (**Figures 4C–4F, and S10**). The inhibitory effect of Leu143Glu and Ile147Glu may be due to the instability of the complex as MlaD(Leu143Glu) and MlaD(Ile147Glu) lost its SDS-resistant hexameric form in the presence of SDS (**Figure S10E**). MlaD’s Tyr152 has been reported to interact with MlaC as shown in a photo-crosslink assay ^50^, suggesting that stable interaction between MlaD and MlaC is important for PL transport. Phe150Glu in the channel retarded the PL transport but did not affect cellular survivability, suggesting its importance in PL transport but not essential(**Figures 4C–4F**). These results are consistent with the known function of MlaD as the mammalian cell entry (MCE) proteins that form hydrophobic channels and transport hydrophobic molecules^45^

**Figure 4.**
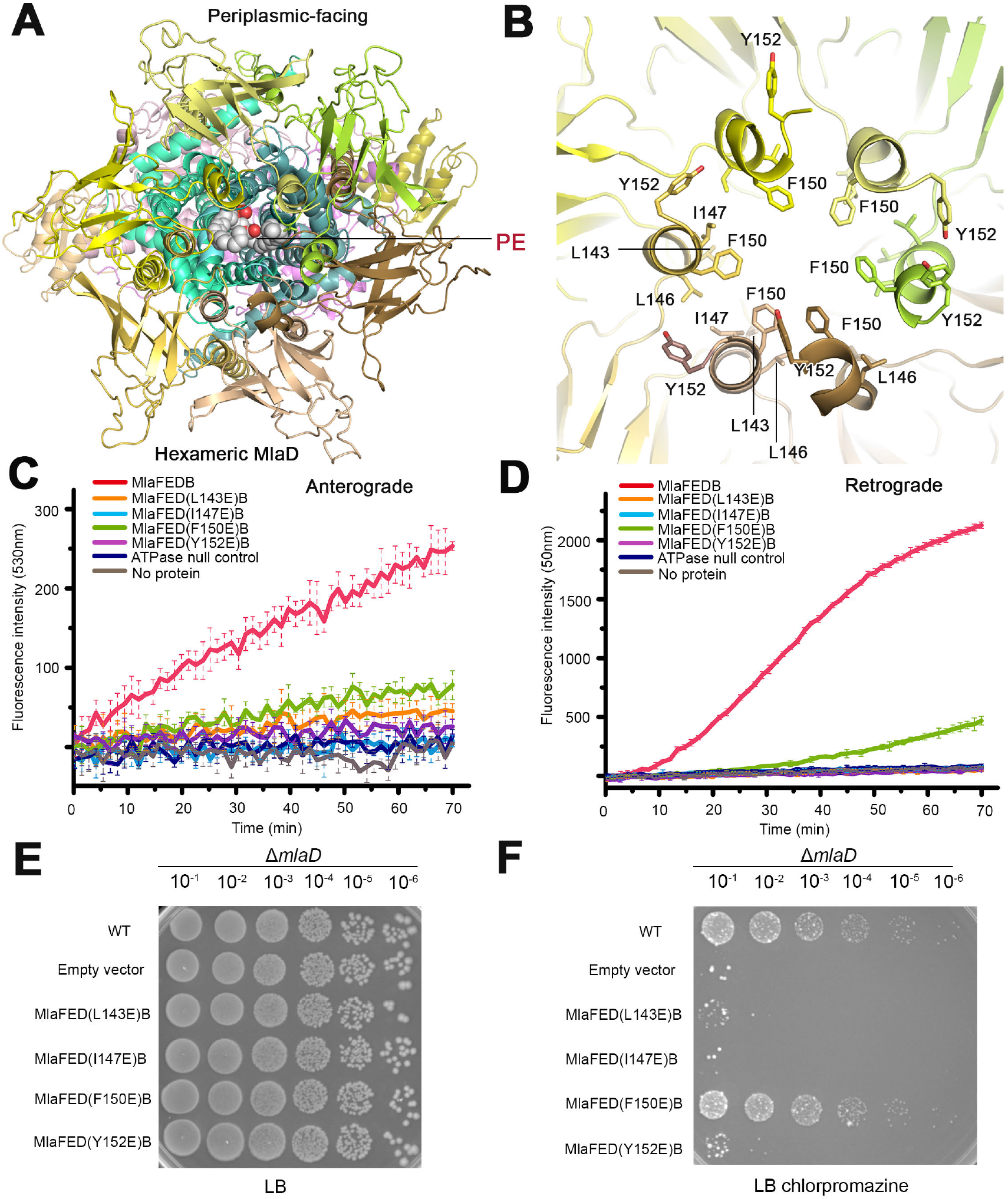
Structural and functional characterization of MlaD. (A) Periplasmic view of MlaFEDB showing the bound PE. (B) The periplasmic central channel of the hexameric MlaD showing the hydrophobic residues L143, I147 and F150 facing inside the channel, and Y152 on the top of the channel. (C and D) FRET assays measuring PL transport in anterograde (C) and retrograde (D) directions in response to mutations of central channel residues. Mutants L143D, I147D and Y152D abolished PL transport, and mutant F150D impaired PL transport. Representative data of n≥3 experiments. (E and F) Cell based assay of MlaD variants using the MlaD deletion strain. The assays were performed on LB plate without (E) or with (F) 120 μg ml^-1^ mM chlorpromazine.

### MlaB stabilizes and regulates MlaFED transporter

MlaB consists of three α-helices and four β-strands, forming a sulphate transfer and anti-sigma-factor antagonist (STAS) domain (**Figure S9D**). MlaB is important for the stability of the complex MlaFEDB. Complex MlaFED cannot be purified without the presence of MlaB^42^. In our cryo-EM structures, the interface area between MlaB and MlaF is 768.4 Å^2^ involving several polar and hydrophobic interactions (**Figures S9E and S9F**). To confirm the importance of these interactions, single MlaB mutants of polar residue Thr52Ala and hydrophobic residues Trp29Glu or Tyr88Glu were generated in the MlaFEDB complex. Amongst these mutants, we failed to purify MlaFEDB(Trp29Glu) or MlaFEDB(Tyr88Glu), suggesting that these hydrophobic residues are crucial for the interaction between MlaB and MlaF thus affecting the stability of the entire complex. Mutant MlaFEDB(Thr52Ala) complex could be purified but completely abolished the ATPase activity of the complex (**Figures S9G and S9H**), which is consistent with a previous report^42^. Furthermore, MlaFEDB(T52A) also failed to transport PL in retrograde direction (**Figures S9I and S9J**). Our structures show that MlaB binds at the a-helical domains of MlaF which in turn interacts with the coupling helices of MlaE (**Figures S9G and S9H**), suggesting that MlaB may play a crucial role in stabilizing the interaction between the NBDs and TMDs of the complex.

### Binding of ADP or AMP-PNP in MlaF

In the ADP- and AMP-PNP-bound structures of MlaFEDB, two extra densities at the two subunits of MlaF are identified as ADP and AMP-PNP respectively with magnesium ions (**Figures 2D, 2E, S9E, and S9F**), which are both interacted by the same set of residues in the active site of the NBDs (**Figures 5A and 5B**). The adenine ring of the nucleotide interacts with Phe16 and Arg18 of MlaF, the ribose interacts with Arg21 and Ile23 (**Figures 5A and 5B**). The third phosphate group of AMP-PNP is surrounded by the positive residues Lys47 and His203 and Mg^2+^ ion which is further coordinated by the negative residue Glu170 (**Figure 5B**). Single mutations Lys47Ala, Glu170Ala or His203Ala severely impaired the ATPase as well as *in vitro* transport activity of the MlaFEDB complex in retrograde direction (**Figures S11A-S11E**), confirming their roles in ATP hydrolysis. The catalytic mutant MlaF(Glu170Ala)EDB showed less effect on anterograde transport of PLs (**Figure S11D**).

**Figure 5.**
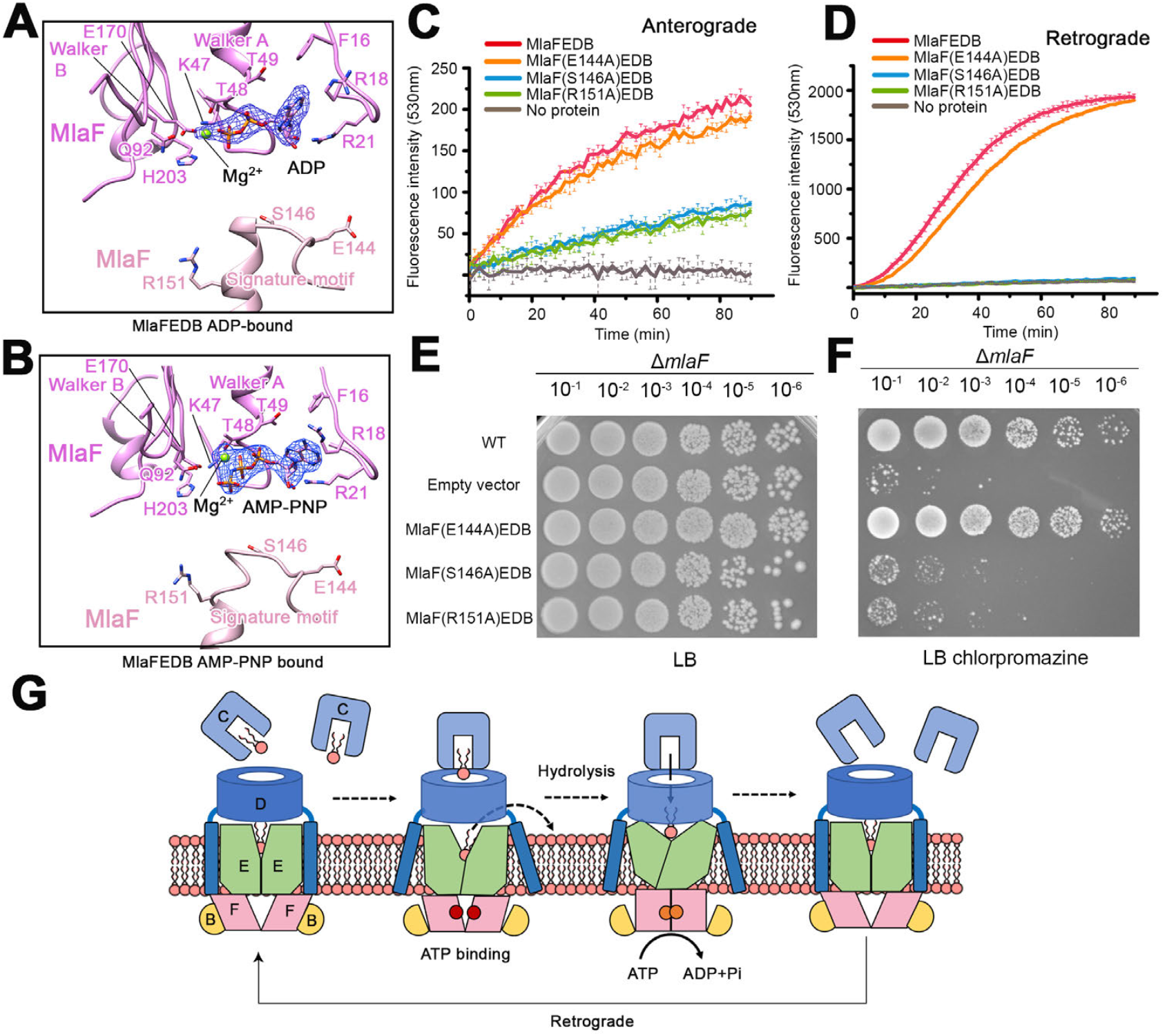
Structural and functional characterization of MlaF. (A and B) ADP (A) or AMP-PNP (B) and Mg^2+^ ions binding residues in the catalytic site of one MlaF subunit and the signature motif of the other MlaF subunit containing conserved residues E144, S146 and R151. (C and D) FRET assays measuring PL transport in anterograde (C) and retrograde (D) directions in response to mutations of MlaF signature motif residues. Representative data of n≥3 experiments. (E and F) Cell based assays of MlaF mutants of signature residues using the MlaF deletion strain. The assays were performed on LB without (E) or with 120 μg ml^-1^ mM chlorpromazine (F). (G) A scheme of a proposed mechanism of the Mla pathway showing MlaE (green), MlaF (pink), MlaB (yellow), MlaD (blue), MlaC (light blue), PL (red). Retrograde transport: MlaC-PL binds to MlaD at the resting state of MlaFEDB. Binding of ATP induce the exit of the PL in the cavity from the last cycle of the retrograde transport. ATP hydrolysis induces dimerization of MlaF and conformational changes of the MlaE to extract PL from MlaD-MlaC bridge into the cavity of MlaE. Hydrolysis products are released and the conformation returns back to the resting state.

MlaF as the NBD of the complex shares folds with other ABC transporters^27,51–53^. MlaF shows 24.54% amino acid sequence identity to LptB, the NBD of LPS transport complex LptBFGC (**Figure S11F**), and structural similarity with an RMSD of 3.81 Å over 358 aligned Cα residues (**Figure S11G**). Unlike LptB, MlaF has an additional C-terminal tail containing residues Gly246 to Ser269, which interacts with the opposite MlaB as well as the opposite unit of MlaF via the residues Phe254, Arg255, Tyr256 and His262 of the C-terminal tail (**Figure S12A**). Single mutations of Tyr256Asp and His262Asp did not reduce ATPase and transport activity of the complex (**Figures S12B-S12E**) or cellular sensitivity to chlorpromazine (**Figure S12F**), suggesting that these C-terminal residues are not involved in the function of the complex. However, we failed to purified MlaFEDB complex with the C-terminal tail being deleted, suggesting that the C-terminal tail of MlaF play a role in stabilizing the complex.

In our current MlaFEDB structure, NBD dimerization is not visualized (**Figure 5B**). It is worth noting that the densities of the bound nucleotides in these MlaFEDB structures are larger than those we obtained for LptBFGC complex ^27^. It may be that the nucleotides in these MlaFEDB structures are not tightly bound in the NBDs, a state before or after dimerization of the NBDs occurred, thus no conformational changes of the complex observed (**Figures 5A and 5B**). We anticipate that the NBD dimerization should be still required for MlaFEDB transporter as other ABC transporters (F**igures S12I-S12M**). Single mutations of the conserved signature motif residues Glu144Ala, Ser146Ala and Arg151Ala of MlaF reduced ATPase activities of the MlaFEDB complexes (**Figures S12B and S12C**). Ser146Ala and Arg151Ala also abolished retrograde PL transport and cellular survivability to chlorpromazine but less affected anterograde FRET signal (**Figures 5C–5F**). Binding of the non-hydrolyzable ADP or AMP-PNP into the MlaFEDB complex did not induce conformational change may also suggest that a conformational change of Mla complex only occurs upon ATP hydrolysis.

It seems that the FRET anterograde transport is not abolished by NBD dimerizing mutations MlaF Ser146Ala and Arg151Ala (**Figures 5C and 5D**), ATPase catalytic mutation MlaF Glu170Ala (**Figures S11D and S11E**), or ATPase activity mutant MlaB Thr52Ala (**Figures S9I and S9J**) but is abolished by absence of ATP (**Figures S1L and S1M**) or ATP binding mutation MlaF Lys47Ala (**Figures 5E and 5F**). These suggest that anterograde transport of PLs is regardless the ATPase activity of MlaFEDB and the retrograde PL transport induced by ATP hydrolysis blocks the path for anterograde (**Figure 1M**). For anterograde PL transport to be observed *in vitro*, it requires an additional cycle of nucleotide binding to expel the bound PL from the cavity, or an apo-MlaC acceptor to passively receive the bound PL anterogradely (**Figures 1B, and S1N**). The continuity of PL transport in either direction is dependent on the PL concentration carried in the MlaC pool.

## Discussion

Our *in vitro* transport assay proposes an ATP-dependent retrograde PL transport from MlaA/OmpF to MlaFEDB and a passive anterograde transport of PL from MlaFEDB to MlaC. MlaFEDB transports PLs bidirectionally *in vitro* but functions predominantly as an ABC retrograde PL transporter. Previously Hughes *et al* observed that apo-MlaC could accept PL from the soluble TM truncated MlaD but not the other way around, proposing an anterograde transport of the PL by the Mla pathway^49^. Our findings show that MlaD is essential for PL transport in both directions (**Figures 4C, 4D, and S10**). Our cryo-EM structures show that the periplasmic MCE channel of MlaD form a continuous transport path with the PL binding cavity of MlaFEB to allow PL trafficking between the IM and periplasm (**Figures. 3A, 3B, and 4B**). The presence of the TM segments of MlaD is important for stabilized complex formation between MlaD and MlaFEB, which allows the coupling of the MCE channel to the ATPase, making energy available for MlaD to overcome the high PLa

ffinity of MlaC to make retrograde transport feasible. Indeed, MlaD shows having a regulatory effect on the ATPase activity of MlaFEB (**Figure 10B**). Functional mutagenesis of MlaFEDB inhibited the *in vitro* retrograde PL transport as well as *in vivo* bacterial survivability to stress, implicating the physiological significance of the Mla system as to retrieve PL from the OM for its asymmetry to maintain bacterial resistance to environmental stress.

Our PL-bound MlaFEDB structure reveals only head-down orientated PE in the cavity of the TMDs (**Figures 2A–2C, 3A, and 3B**). Mutation of the PE polar head binding residue Glu98Arg severely impairs the retrograde transport of PL and cellular survivability to stress, suggesting that the head-down orientated PL observed in our cryo-EM structure is being transported retrogradely (**Figures 3F-H**). Interestingly, Glu98Arg showed no inhibitory effect to the anterograde transport, suggesting that the PL transported for anterograde may not necessarily be in the same orientation as the retrograde (**Figure 3F**). The inverted detergent molecules bound in the structures may suggest that PL could also be bound in a head-up orientation with the acyl chains interacting with the hydrophobic residues in the upper cavity. MlaFEDB extracts PL from proteoliposomes and passes it to apo-MlaC independent of ATP (**Figure 1C**), suggesting that the hydrophobic force in the substrate-binding cavity is sufficient for PL extraction from the IM yet allows PL diffusion through the MCE of MlaD to an interacted apo-MlaC along the gradient of PL affinity^54^. In the case that MlaC is already in a PL bound state, anterograde transport is blocked and the retrograde PL transport is prioritized with the action of ATPase is involved. We showed the tendency of PL (from IM proteoliposomes)-bound MlaC to associate with the OM proteoliposomes but no evidence that anterograde PL transport involves MlaA/OmpF (**Figures S1O and S1P**). However, the bidirectional transport activity of MlaFEDB *in vitro* and the published IM accumulation of PLs in Mla mutants may imply some physiological relevance of the anterograde transport, probably for supporting bacterial cell growth^48^. Other PL transport pathways YebST^55,56^ and PqiABC^45,57^ may act synergistically with the Mla system for anterograde transport of PLs but evidence needed to prove. Nevertheless, the finding of prioritized retrograde PL transport is reasonable as urgent bacterial response for survival in environmental stress.

The structural characteristics of MlaFEDB requiring several extra stabilizing regulatory factors comparing to other canonical ABC transporters may provide extra constraints for the stability of the complex, forcing the complex to rapidly return to its rested conformation after each cycle of ATP hydrolysis. The current resting conformation shows an outward-opening cavity, which accommodates a PL transported from the last cycle of the retrograde transport. We speculate that the bound “head-down” PL only exits into the IM when ATP is bound and the simultaneous ATP hydrolysis induces conformational changes to extract next PL from the MlaD-MlaC-PL bridge into the cavity. The continuity of the PL retrograde transport is dependent on the presence of ATP and PL concentration in the MlaC pool charged by MlaA/OmpF from the OM (**Figure 5F**).

## Methods

### Expression and purification of MlaFEDB complex

Genes encode for *mlaFEDCB* of *E. coli* K-12 strain was chemically synthesized and cloned into a pTRC99a plasmid using EcoRI and KpnI restriction enzymes, resulting a *pTRC99a-mlaFEDCB* construct with an octa-histidine (8 × His) tag at the C-terminus of MlaB. The resulting plasmid was transformed into *E. coli C43 (DE3)* cells (Novagen) for protein expression. The bacterial cells were grown in Luria broth (LB) supplemented with 100 μg ml^-1^ ampicillin at 37 °C until the optical density of the culture reached 0.8 at a wavelength of 600nm (OD_600nm_). Protein expression was induced with 0.1 mM isopropyl-β-D-thiogalactopyranoside (IPTG) at 20 °C for 12h.

Cultures were harvested by centrifugation and the cell pellets were resuspended in lysis buffer containing 20 mM Tris-Cl, pH 7.8, 300 mM NaCl, 10% (v/v) glycerol and supplemented with 0.1 mM phenylmethylsulphonyl fluoride (PMSF). The cells were lysed by three passes through a cell disrupter and cell debris was removed by centrifugation at 18,000 × *g* for 15 min at 4°C. Membranes were pelleted by ultracentrifugation at 100,000 × *g* for 1h at 4 °C and solubilized with 20 mM Tris-Cl, pH 7.8, 300 mM NaCl, 10% (v/v) glycerol, 10 mM imidazole and 1% (w/v) *n*-dodecyl-β-D-maltopyranoside (DDM; Anatrace) by stirring at room temperature for 1h. The suspension of solubilized protein was ultracentrifuged at 100,000 × *g* for 1h before being loaded onto a 5 ml HisTrap HP column (GE HealthCare). The column was washed with buffer containing 20 mM Tris-Cl, pH 7.8, 300 mM NaCl, 10% (v/v) glycerol, 50 mM imidazole and 0.05% (w/v) Lauryl Maltose Neopentyl Glycol (LMNG; Anatrace), then bound protein was eluted with 20 mM Tris-Cl, pH 7.8, 300 mM NaCl, 10% (v/v) glycerol, 300 mM imidazole and 0.05% LMNG. The protein eluted from Histrap HP column was further purified by size-exclusion chromatography using a Superdex 200 Increase 10/300 column (GE Healthcare) equilibrated in 20 mM Tris-Cl, pH 7.8, 150 mM NaCl and 0.05% LMNG. The protein fractions of MlaFEDB complex with the best purity were obtained for cryo-sample preparation. MlaC was not detected as part of the complex by SDS-PAGE (**Figure S1A**). For nucleotides-bound complexes, the purified MlaFEDB protein complex was incubated with 5 mM β-γ-imidoadenosine 5’-phosphate (AMP-PNP) or 5 mM ADP and 2 mM MgCl_2_ for 1h at room temperature before cryo-sample preparation. To obtain substrate-bound complex, the purified protein was incubated with 5 mM E. coli polar lipid extract (Avanti Polar Lipids) at room temperature for 1h.

### Expression and purification of MlaC

Gene fragment encoding MlaC (residues 21-211) from *E. coli* K-12 strain with the signal peptide removed was amplified and cloned into a modified pET28a plasmid containing an octa-histidine (8 × His) tag and a SUMO-fusion followed by a tobacco etch virus (TEV) protease cleavage site (8 × His-SUMO-TEV-MalC21-211). The recombinant plasmid was transformed into *E. coli C43 (DE3)* cells for protein expressions. The bacterial cells were grown in LB supplemented with 50 μg ml^-1^ Kanamycin at 37 °C until the optical density of the culture reached 0.8 at a wavelength of 600_nm_. Protein expression was induced by addition of 0.1 mM IPTG and incubated for 12h at 20 °C. The cells were harvested by centrifugation at 5,000 × g for 20 min, resuspended in buffer containing 20 mM Tris-Cl, pH 7.8, 10% (v/v) glycerol, 500 mM NaCl, supplemented with 1 mM DNase (Sigma) and 1 mM PMSF. The cells were lysed using a cell disruption and the cell debris was removed by centrifugation at 120,000 × g for 25 min at 4 °C. The supernatant was then loaded onto a Ni-NTA column (Qiagen) and the column was washed with 20 mM Tris-Cl, pH 7.8, 10% (v/v) glycerol, 500 mM NaCl and 30 mM imidazole. The protein was eluted with 20 mM Tris-Cl, pH 7.8, 10% (v/v) glycerol, 500 mM NaCl and 300 mM imidazole. The excessive imidazole was removed using a desalting column (Hi-Prep^™^ 26/10, GE Healthcare). The 8 × His-SUMO was cleaved by TEV proteinase and the cleaved fragment was removed through the affinity of the His-tag to a Ni-NTA column. The protein was further purified using size-exclusion chromatography with buffer containing 20 mM Tris-Cl, pH 7.8 and 150 mM NaCl and the fractions were examined by SDS-PAGE (**Figure S9H)**. Protein fractions with the highest purity of MlaC21-211 were pooled. The protein was concentrated to 7mg/ml for the assays.

### Expression and purification of MlaD and MlaFEB

Gene fragments encoding full-length MlaD or MlaD with the transmembrane (TM) a-helix removed (hereafter MlaDΔTM) from *E. coli* strain K-12 were amplified and cloned into plasmid pET28a with a BamH I and Hind III restriction sites respectively. The resulting plasmid contain an octa-histidine (8 × His) tag and a SUMO-fusion followed by a tobacco etch virus (TEV) protease cleavage site upstream of MlaD or MlaDΔTM, and a strep-tag at c-terminal (8 × His_SUMO_TEV_MlaD_Strep-tag or 8 × His_SUMO_TEV_MlaDΔTM_Strep-tag). The recombinant plasmids were transformed into *E. coli C43 (DE3)* cells respectively for protein expressions. The soluble Strep-tagged MlaDΔTM protein was overexpressed and purified using the same protocol as described for MlaC above. The 8 × His-SUMO was cleaved by TEV proteinase and the cleaved fragment was removed through the affinity of the His-tag to a Ni-NTA column. The protein was further purified using size-exclusion chromatography with buffer containing 20 mM Tris-Cl, pH 7.8 and 150 mM NaCl and the fractions were examined by SDS-PAGE (**Figure S10A**).

To overexpress the full-length Strep-tagged MlaD, the bacterial cells were grown in LB supplemented with 50 μg ml^-1^ Kanamycin at 37 °C until the optical density of the culture reached 0.8 at a wavelength of 600nm. Protein expression was induced by addition of 0.1 mM IPTG and incubated for 12h at 16 °C. The cells were harvested by centrifugation at 5000 × g for 20 min, resuspended in buffer containing 20 mM Tris-Cl, pH 7.8, 10% (v/v) glycerol, 500 mM NaCl, supplemented with 1 mM PMSF. The cells were lysed by three passes through a cell disrupter and cell debris was removed by centrifugation at 18,000 × *g* for 15 min at 4°C. Membranes were pelleted by ultracentrifugation at 100,000 × *g* for 1h at 4 °C and solubilized with 20 mM Tris-Cl, pH 7.8, 300 mM NaCl, 10% (v/v) glycerol, 10 mM imidazole and 1% (w/v) DDM by stirring at room temperature for 1h. The suspension of solubilized protein was ultracentrifuged at 100,000 × *g* for 1h before being loaded onto a 5 ml HisTrap HP column. The column was washed with buffer containing 20 mM Tris-Cl, pH 7.8, 300 mM NaCl, 10% (v/v) glycerol, 50 mM imidazole and 0.05% (w/v) DDM, the bound protein was eluted with 20 mM Tris-Cl, pH 7.8, 300 mM NaCl, 10% (v/v) glycerol, 300 mM imidazole and 0.05% DDM. The protein buffer was exchanged to 20 mM Tris-Cl, pH 7.8, 300 mM NaCl, 10% glycerol, 10 mM imidazole and 0.05% DDM using a desalting column. The 8 × His-SUMO was cleaved by TEV proteinase and the cleaved fragment was removed through a Ni-NTA column. The protein was further purified using size exclusion chromatography in 20 mM Tris-Cl, pH 7.8 and 150 mM NaCl and 0.05% DDM and examined by SDS-PAGE. Protein fractions with the highest purity of Strep-tagged MalD were pooled (**Figure S10A**).

To overexpress MlaFEB, the pTRC99a-*mlaFEDCB* construct with an octa-histidine (8 × His) tag at the C-terminus of MlaB was used as a template to clone the gene for MlaFEB by PCR. The MlaFEB protein complex was expressed and purified using the same protocol as described for MlaFEDB above (**Figure S10A**).

The purified Strep-tagged full length MlaD or MlaDΔTM form hexamer *in vitro* (**Figure S10A**), which were individually incubated with the purified His-tagged MlaFEB (hereafter MlaFEB+MlaD or MlaFEB+MlaD(ΔTM)) with molar ratio of 1:1 at room temperature for 6h to allow complex formation *in vitro*. The incubated mixture was first loaded onto a NI-NTA column, which was washed with buffer containing 20 mM Tris-Cl, pH 7.8, 300 mM NaCl, 10% (v/v) glycerol, 10 mM imidazole and 0.05% (w/v) DDM, and eluted with 20 mM Tris-Cl, pH 7.8, 300 mM NaCl, 10% (v/v) glycerol, 300 mM imidazole and 0.05% DDM. SDS-PAGE show that his-tagged MlaFEB could only pull-down full length MlaD but not MlaDΔTM (**Figure S10A**).

The eluted samples containing His/Strep-tagged MlaFEB+MlaD was further purified by the Strep column, which was washed with buffer containing 20 mM Tris-Cl, pH 7.8, 300 mM NaCl, 10% (v/v) glycerol, and 0.05% (w/v) DDM, and the bound protein was eluted with 20 mM Tris-Cl, pH 7.8, 300 mM NaCl, 10% (v/v) glycerol, 3 mM d-Desthibiotin (Sigma) and 0.05% DDM. The protein buffer of the eluted samples was exchanged to 20 mM Tris-Cl, pH 7.8, 300 mM NaCl, 10% glycerol, 10 mM imidazole and 0.05% DDM using a desalting column. The purified protein of MlaFEB+MlaD complex was examined by SDS-PAGE (**Figure S10A**).

### Expression and purification of MlaA/OmpF complex

Gene fragment encoding MlaA from *E. coli* K-12 strain was amplified and cloned into a pTRC99a plasmid using EcoRI and KpnI restriction enzymes, resulting a pTRC99a-*mlaA* construct with an octahistidine (8 × His) tag at the C-terminus of MlaA. The recombinant plasmid was transformed into *E. coli C43 (DE3)* cells for protein expressions. The transformed cells were grown in LB supplemented with100 μg ml^-1^ ampicillin at 37 °C until the optical density of the culture reached 0.85 at 600nm. Protein expression was induced by addition of 0.1 mM IPTG and incubated for 4h at 15 °C. The cells were harvested by centrifugation at 5,000 × g for 20 min, resuspended in buffer containing 20 mM Tris-Cl, pH 7.8, 10% (v/v) glycerol, 500 mM NaCl, supplemented with 1 mM DNase and 1 mM PMSF. The cells were lysed by three passes through cell disrupter and cell debris was removed by centrifugation at 18,000 × *g* for 15 min at 4°C. Membranes were pelleted by ultracentrifugation at 100,000 × *g* for 1h at 4 °C and solubilized with 20 mM Tris-Cl, pH 7.8, 300 mM NaCl, 10% (v/v) glycerol, 10 mM imidazole and 1% (w/v) DDM by stirring at room temperature for 1h. The suspension of solubilized protein was ultracentrifuged at 100,000 × *g* for 1h before being loaded onto a 5 ml HisTrap HP column. The column was washed with buffer containing 20 mM Tris-Cl, pH 7.8, 300 mM NaCl, 10% (v/v) glycerol, 50 mM imidazole and 0.05% (w/v) DDM, then bound protein was eluted with 20 mM Tris-Cl, pH 7.8, 300 mM NaCl, 10% (v/v) glycerol, 300 mM imidazole and 0.05% DDM. The protein was further purified using size exclusion chromatography with 20 mM Tris-Cl, pH 7.8 and 150 mM NaCl and 0.05% DDM. Fractions containing the target proteins of MlaA/OmpF complex were examined by SDS-PAGE (**Figure S9H**).

### Reconstitution of MlaFEDB and MlaA/OmpF complex into proteoliposomes

An *E. coli* polar lipid (Avanti Polar Lipids) was solubilized in chloroform, then dried with nitrogen gas to form a lipid film. The lipid film was resuspended at a concentration of 10 mg ml^-1^ in buffer containing 20mM Tris, pH 7.8, and 150 mM NaCl, and subjected to sonication, and then passed through a polycarbonate membrane with 0.4 μm pore size to generate liposomes. Liposomes were destabilized by adding 1.6 mmol DDM per mg lipid, followed by incubation with MlaFEDB or relevant mutations or MlaA/OmpF at a molar ratio of 20:1 for 1h on ice. To remove the detergents and to incorporate protein into the liposomes, the solution was incubated with 120 mg Bio-Beads SM2 (Bio-Rad). After a final 3h incubation with Bio-Beads at 4°C, proteoliposomes −1 were collected and resuspended at 1 mg ml^−1^ in 20 mM Tris, pH 7.8, and 100 mM NaCl for ATPase activity assay.

1-palmitoyl-2-oleoyl-glycero-3-phosphocholine (POPC) lipid film (Avanti Polar Lipids) was resuspended at a concentration of 10mg/ml in buffer 20mM Tris, pH 7.8, 150 mM NaCl, 1mM ATP and 2mM MgCl_2_ and subjected to sonication, and then passed through a polycarbonate membrane with 0.4μm pore size to generate liposomes. Liposomes were destabilized by adding 1.6 mmol DDM per mg lipid, followed by incubation with MlaFEDB or MlaA/OmpF at a ratio of 20:1 on ice for 1h. For detergents removal, Bio-beads SM2 were added 120 mg into the liposomes and incubated for 3 h at 4°C and proteoliposomes were collected for thin-layer chromatography assay.

### In vitro phospholipid (PL) transport assay by thin-layer chromatography (TLC)

A *pTRC99a-mlaFEDCB* construct with an octahistidine (8 × His) tag at the C-terminus of MlaB or a pTRC99a-*mlaA* construct with an octa-histidine (8 × His) tag at the C-terminus of MlaA was further cloned with a TEV protease cleavage site before the His-tag of MlaB or MlaA. When necessary, the C-terminal His-tag from MlaB or MlaA was cleaved by TEV proteinase and the cleaved fragment was removed by a Ni-NTA affinity column.

To test the transport direction of PLs of the Mla system, we also detected PL transported to apo-MlaC by TLC. Purified MlaC contains endogenous PL, which needs to be removed before the transport assay. In some TLC experiments, we also used apo-MlaFEDB and apo-MlaA/OmpF to construct proteoliposomes. To obtain apo-proteins, Ni-NTA column bound His-tagged protein was washed with 40 columns volume of extraction buffer containing 50 mM Tris pH 8.0, 300 mM NaCl, 10 mM imidazole and 1% DDM, followed by 40 columns volume of elution buffer containing 50 mM Tris pH 8.0, 300 mM NaCl, 10 mM imidazole and 0.05% DDM. The resulted apo-protein was further purified using size exclusion chromatography in the gel filtration buffer of 20 mM Tris-Cl, pH 7.8 and 150 mM NaCl without (for MlaC) or with (for MlaFEDB or MlaA/OmpF) 0.05% DDM. For TLC experiment, we used un-labeled E. coli polar lipids or POPC to construct MlaFEDB or MlaA/OmpF proteoliposomes as described before to differentiate the origin of the transported PL. Apo-MlaC used in the reaction was in excess 20mg for every 1mg of the proteoliposome mix (MlaFEDB:MlaA/OmpF=1:1). The transport reaction was allowed for 30min in the presence or absence of 1mM of ATP and 2mM MgCl_2_. After reaction, proteoliposomes were collected by centrifugation at 300,000 g for 45 min, and the supernatant (MlaC) was subjected to size exclusion chromatography using a Superdex 200 Increase 10/300column, and the fractions containing MlaC was collected. Methanol and chloroform was used to extract PLs from MlaC (protein samples : methanol :chloroform =2:2:1v/v/v) according to the described method^49^. The lipids were re-dissolved in 200 μl of chloroform and 5 μl sample was loaded onto a Silica TLC plate (Silica gel 60 F254–Merck Millipore) and separated of lipids by solvent system (chloroform: methanol: water=65:25:4, v/v/v). The TLC plate was dried for 30 min and stained with 10% (w/v) phosphomolybdic acid in ethanol and heated until staining could visible.

### In vitro phospholipid transport assay by Fluorescence Resonance Energy Transfer (FRET)

To monitor lipid transfer between liposomes via MlaFEDB and MlaA/OmpF, we adopted fluorescent de-quenching assays as described previously ^58^. Fluorescent labelled PE was used for constructing the donor proteoliposomes and unlabelled POPC was used for acceptor proteoliposomes. For fluorescent (donor) proteoliposomes, *E. coli* polar lipid extract, 1,2-dipalmitoyl-*sn*-glycero-3-phosphoethanolamine-*N*-(7-nitro-2-1,3-benzoxadiazol-4-*yl*) (NB-PE) and 1,2-dioleoyl-*sn*-glycero-3-phosphoethanolamine-*N*-(lissamine rhodamine B sulfonyl) (Rhod-PE) (Avanti Polar Lipids) lipid film were respectively resuspended at 10 mg ml^-1^, 1 mg ml^-1^ and 1 mg ml^-1^ in buffer containing 20mM Tris, pH 7.8, 150 mM NaCl, 1mM ATP and 2mM MgCl_2_. The *E. coli* polar lipid extract: NB-PE: Rhod-PE at a ratio of 92:6:2 (w/w) were mixed and subsequently sonicated to homogenize and passed through a polycarbonate membrane with 0.4 μm pore size to generate liposomes. Liposomes were destabilized by adding 1.6 mmol DDM per mg lipid, followed by incubation with MlaFEDB or MlaA/OmpF at a ratio of 20:1 on ice for 1h. For detergents removal, Biobeads SM2 were added 120 mg into the liposomes and incubated for 3 h at 4°C and proteoliposomes were collected for FRET assay. The unlabelled POPC proteoliposomes were constructed same as before.

MlaFEDB and MlaA/OmpF complexes were respectively reconstituted into labelled or unlabeled proteoliposomes at molar ratio of 1:1, and then 10μl of each reaction was diluted into 130μl buffer containing 20mM Tris, pH7.8, 150 mM NaCl, 1mM ATP and 2mM MgCl_2_. The mixture was divided into two groups: one group was added with 20mg MlaC per mg of MlaFEDB (the test group) and the other group added with same volume of buffer (the control group). 20μl samples from either group were loaded onto each well of a 384-well plate. Transported NB-PE or rhod-PE from the labelled proteoliposomes (donor) to the unlabelled proteoliposomes (acceptor) show increased de-quenched NB fluorescence (530nm) or decreased FRET excited rhod (590nm), which was monitored using a Biotek Cytation 3 Cell Imaging multi-mode reader every 10s until 60 min reaction. Because of the initial delay until the first record of the transport, the NB fluorescent at the time point zero varies depending on the transport activities in the reaction. Quenched NB fluorescence in the donor proteoliposomes was set to 0%.

Each PL transport assay has two additional controls, one is used empty liposome without protein complex (thereafter no protein) as a control and other is ATPase null control using mutant MlaEDBFK47A. For *in vitro* formed complex, purified protein MlaFEB+MlaD was constructed for the IM proteoliposomes for the PL transport assay. To test the function of MlaD(ΔTM), MlaFEB was constructed for IM proteoliposomes and soluble MlaD(ΔTM) was added into the PL transport assay. All assays have done in triplicates and repeated at least three times.

### Cellular survivability assay to chlorpromazine

All single mutations were generated following the site-directed mutagenesis protocol as previously described^59^. The null strains of *MlaD* (JW3160)*, MlaE* (JW3161)*, or MlaF* (JW3162) used for functional assays were from the Keio collection^60^, which were conducted by P1 transduction of kanamycin resistance. The *pTRC99a-MlaFEDB* construct with His8-tag at the C terminus of MlaB was used as a template by PCR for the *E. coli* MlaFEDB mutagenesis.

These single mutants of MlaD or MlaE or MlaF were transformed into the appropriate deletion stain. The transformed *E. coli* cells were grown on LB agar plate supplemented with antibiotics (kanamycin 50 μg ml^-1^, ampicillin 100 μg ml^-1^) at 37 °C for 16h. Single colonies of each transformation were inoculated into 10 ml LB medium supplemented with the antibiotics. The cells were cultured in an incubator at 200 r.p.m. at 37 °C for 12 h and the sub-cultured cells were used for the functional assays. The empty plasmid pTRC99a-amp was used as the negative control, while the plasmid pTRC99a-*MlaFEDB* was used as the positive control. Cell pellets were harvested, washed twice and diluted in sterile LB medium to the OD600 of 0.5. Cell viability assay with ten-fold serial dilution were performed. The dilution range was from 10^-1^ to 10^-6^ and 5μl of the diluted cells was dripped onto the LB agar plates containing 50 μg ml^-1^ kanamycin and 100 μg ml^-1^ampicillin with or without 120 μg ml^-1^ chlorpromazine. Cell growth was observed after overnight culture at 37 °C. All the assays were performed in triplicate and mutations with leak-expression were purified using same methods as the MlaFEDB described as the above.

### ATPase activity assay

ATPase activity assay was performed using ATPase/GTPase Activity Assay Kit (Bioassay systems) as previously reported^27^. The phosphate standards and blank control for colorimetric detection was prepared according to the manufacturer’s instructions (ATPase/GTPase Assay Kit, Bioassay systems). An aliquot of 1μl (1-2 mg ml^-1^) of purified MlaFEDB or relevant mutations in DDM or in proteoliposomes was mixed with 4μl buffer containing 20 mM Tris, pH 7.8, 150 mM NaCl, followed by adding 5μl assay buffer (ATPase/ GTPase Assay Kit). The mixtures were added to 30μl reaction solution with 20μl assay buffer plus 10μl 4 mM ATP solution and incubated at room temperature for 15 min. The reaction was terminated by adding 200μl reagent (ATPase/ GTPase Assay Kit) into each sample and further incubated for 30 min at room temperature. The absorbance at 600nm was measured. For AMP-PNP or ADP inhibition, MlaFEDB was incubated with AMP-PNP or ADP concentration varying from 0 to 3mM and 5 mM MgCl_2_ at 37°C for 30 min before ATPase activity assay was performed. ATPase activities of all samples were determined using the mean value of the samples according to the linear regression of standards. All experiments were repeated at least three times.

### Electron microscope sample preparation and data acquisition

2.5μl of purified protein complex at a concentration of approximately 4mg ml^-1^ was applied to glow-discharged Quantifoil holey carbon grids (R1.2/1.3, 300 mesh, copper). Grids were blotted for 3.5s with the environmental chamber set at 95% humidity and flash-frozen in liquid ethane cooled by liquid nitrogen using Vitrobot Mark IV (FEI). Grids were imaged with a Titan Krios (FEI) electron microscope, operated at 300 keV equipped with a K2 Summit electron counting direct detection camera (Gatan). All cryo-EM images were recorded in super-resolution mode using the automated data collection program SerialEM^61^ and images were acquired at a nominal magnification of 29,000×, corresponding to a calibrated physical pixel size of 1.014 Å. Defocus range was set between −1.5 to −2.5 μm. Each image was collected at an exposure time of 8 s and dose-fractionated to 40 frames with a dose rate of about 7 counts per second per physical pixel.

### Image processing

The beam-induced motion correction of image stacks were carried out using MotionCor2^62^ to generate 2x binned average micrographs and dose-weighted micrographs with a pixel size of 1.014 Å. The contrast transfer function parameters of these average micrographs were estimated by Gctf^63^. Other procedures of data processing were performed in RELION^64^. For the ADP-bound MlaFEDB structure, 2,364,885 particles were automatically selected, and finally 194,246 particles were selected for 3D refinement and a reported 3.72 Å resolution map was generated after post-processing with a B-factor of −182 Å^2^. The data processing details are summarized in Figure S4.

The data processing procedures of phospholipid-bound MlaFEDB were similar to the procedures of ADP-bound MlaFEDB. 1,910,766 particles were automatically selected, and then two-dimensional (2D) and three-dimensional (3D) classifications were carried out to select consistent particle classes. Finally, 97,672 particles were selected for 3D refinement and a reported 3.1 Å resolution map was generated after post-processing with a B-factor of −77 Å^2^. The data processing details are summarized in Figure S3.

The data processing procedures of AMP-PNP bound MlaFEDB were similar to the procedures of ADP-bound MlaFEDB. 793,723 particles were automatically selected, and then two-dimensional (2D) and three-dimensional (3D) classifications were performed to select consistent particle classes. Finally, 340,974 particles were selected for 3D refinement and a reported 3.4 Å resolution map was generated after post-processing with a B-factor of −172 Å^2^. The data processing details are summarized in Figure S5.

The data processing procedures of apo MlaFEDB were similar to the procedures of MlaFEDB AMP-PNP bound. 1,177,885 particles were automatically selected, and then two-dimensional (2D) and three-dimensional (3D) classifications were performed to select consistent particle classes. Finally, 76,541 particles were selected for 3D refinement and a reported 3.3 Å resolution map was generated after post-processing with a B-factor of −85 Å^2^. The data processing details are summarized in Figure S2.

### Model building and refinement

The crystal structure of the periplasmic domain of MlaD from *E. coli* (PDB code: 5UW2) was fitted into the cryo-EM map of ADP bound MlaFEDB at 3.72 Å using UCSF Chimera^65^. The dimeric MlaE were built using Phenix automatically^66^. The model of MlaF and MlaB was generated using Phyre2 server^67^ and fitted into the cryo-EM densities using COOT^68^. Four TM helices of MlaD were built manually in COOT, while the densities of other two TM helices interacted with the N-terminal residues of MlaE were not clear to build. The high resolution cryo-EM maps allow the side chain assignments (Figure S4). The densities of the loop between the TM helix (residues A29-E37) and the periplasmic domain (residues E119 to A126) of MlaD, as well as the C-terminal tail (residues Y152-K183) of MlaD were not clearly observed, suggesting they are flexible. There is a density at the cavity of MlaFEDB for detergent LMNG used for the protein purification, but only one sugar was modelled in the head group of LMNG. There are two additional densities at dimeric MlaF molecules for ADP, which is bound at the catalytic site of the nucleotide binding domains. The LMNG and ADP molecules are fitted manually in the densities.

The structure of ADP-bound MlaFEDB was initially manually adjusted and fitted into the cryo-EM map of apo MlaFEDB with the resolution of 3.3 Å using UCSF Chimera^65^ (**Figure S2**). There are two densities were identified in the cavity of the native MlaFEDB as LMNG molecules (**Figure S2F**). The bottom one is superimposable to that of ADP-bound MlaFEDB, but the top one rotates almost 90 degree in anticlockwise direction along y-axis relative to the bottom LMNG (**Figures S8A-S8C**). Both head groups of LMNG point to the periplasm (**Figure S8G**). Densities for all six TM helices of hexameric MlaD were clear, and six TM helices of MlaD were manually built in the structure. The resolution of the periplasmic domain of MlaD is in the range of 4.0-5.0 Å, where the periplasmic domain of MlaD was docked into the density map of native MlaFEDB.

The apo MlaFEDB structure was manually adjusted and fit into the PL-bound MlaFEDB cryo-EM map with the resolution of 3.1 Å in COOT (**Figure S3**). There is a density at the cavity, which was identified as a PE molecule. The head group of PE points to the cytoplasmic side, while the two tails of PE point to the periplasmic side. PE molecule is occupied in the same position as the that of LMNG in native MlaFEDB. However, the orientation is different from that of the LMNG in the native MlaFEDB. Map densities for TM helixes of MlaD were clear, and six TM helixes were built in the map. Density for hexameric periplasmic domains was in low resolution at 4.5-5 Å, where the hexameric periplasmic domains were docked in the map density.

The ADP-bound MlaFEDB structure was fitted into the AMP-PNP bound MlaFEDB cryo-EM map with the overall resolution of 3.5 Å manually using COOT (**Figure S5**). AMP-PNP densities were identified at the active site of MlaF, and two AMP-PNP molecules were added into the map using COOT. However, the densities of two of the TM helices of MlaD that are also not visible in the AMP-PNP-bound MlaFEDB structure, thus four TM helices of MlaD were built in the model.

All the structures were refined using PHENIX^66^ with phniex.real_space_refine, and the statistics of the refinement are listed in Table S1.

## Author Contributions

H.D. and X.T. conceived and designed the experiments. X.T. and H. D. made the constructs for protein expression. X.T., W. Q., Q. L. and K. Z. expressed, purified the proteins. Y. C., W. Q., Q. L., T. W., K. Z., Z. Z., C. Z., J.C., X.W. and X. Z. did the mutagenesis, ATPase activities, TLC and the transport assays. X.T., H.D., W. Q. Q. L. and K. Z. prepared the samples. S. C., Z. J. and X. Z. undertook data collection, processed electron microscope data and structure constitution. H. D., X. T. and C.D. did the model building and refinement. H. D., X. T. wrote the manuscript and X. Z., S. C. and C.D. revised the manuscript.

## Acknowledgements

We thank Y. Wei and B. Dong for supporting the project and Y. Zhang for advice in cryo-sample preparation. This Work was supported by grants from the National Key Research and Development Program of China (2017YFA0504803, 2018YFA0507700) and the Fundamental Research Funds for the Central Universities (2018XZZX001-13) to X.Z.; The National Natural Science Foundation of China (31900039 to X. D. and 81971974 to H. D.); The Fundamental Research Funds for the central Universities and supported by National Clinical Research Centre for Geriatrics, West China Hospital (Z2018B01) to H. D.; The National Key Research and Development Program of China (2017YFC0840100, 2017YFC00840101); Laboratory and equipment management, Zhejiang University (SJS201814) to S. C. and Wellcome Trust investigator award (WT106121MA) to C.D.

