## Supplementary file for "Structural insight into outer membrane asymmetry maintenance of Gram-negative bacteria by the phospholipid transporter MlaFEDB"

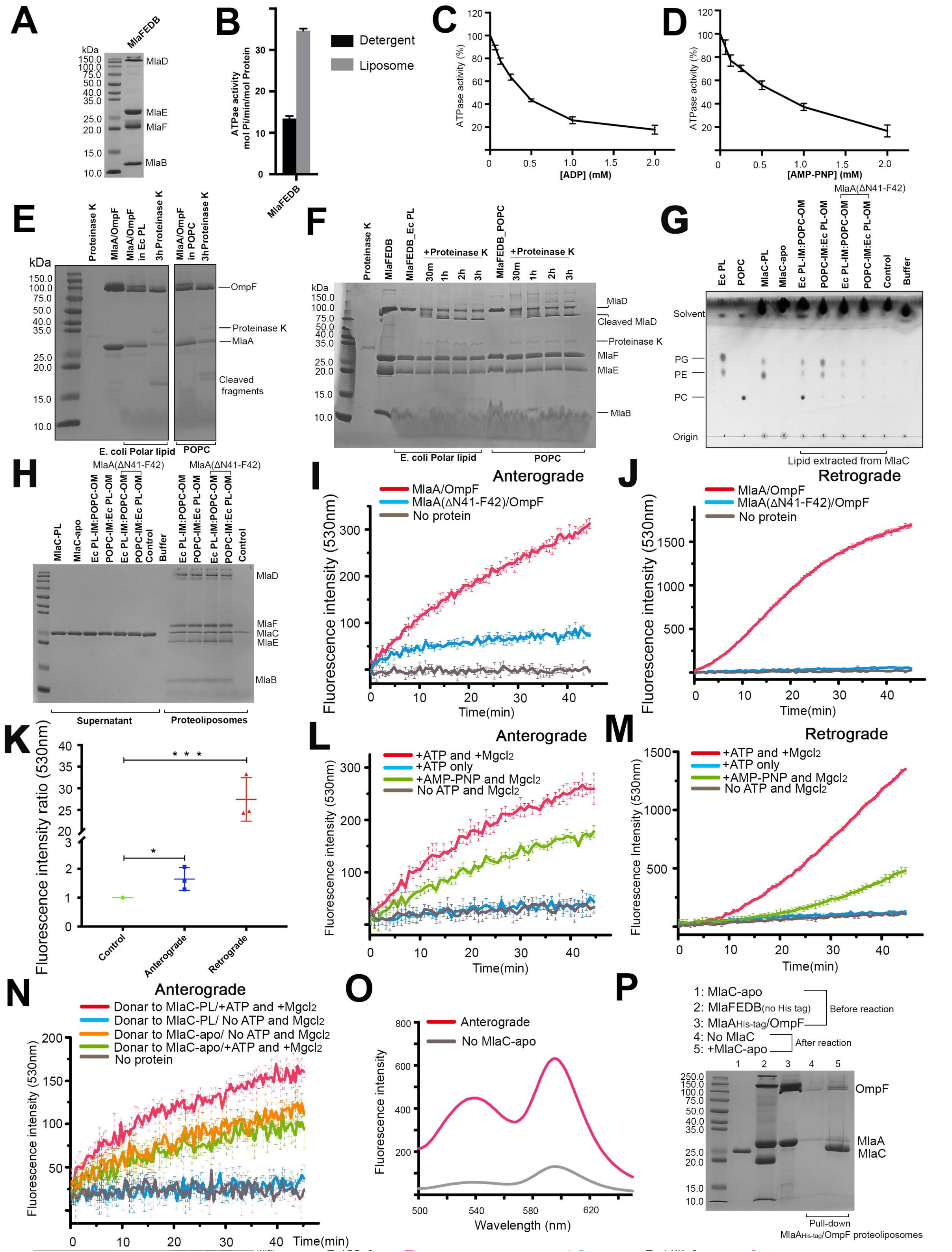

**Figure S1. The setup of the proteoliposomes-based transport system limited proteolysis with proteinase K shows that proteins orientation and *in vitro* PL transport**. (A) Coomassie brilliant blue staining of the purified MlaFEDB on SDS-PAGE. (B) The relative ATPase activities of purified MlaFEDB. MlaFEDB shows higher ATPase activities in liposomes than in detergent. (C and D) The relative ATPase activities of MlaFEDB in the presence of ADP (C) or AMP-PNP (D). (E and F) SDS-PAGE analysis of MlaA/OmpF (E) or MlaFEDB (F) constitution ratio and orientation ratio in proteoliposomes of E. coli polar lipids or POPC with or without proteinase K treatment. (G) Thin-layer chromatogram (TLC) showing transported PE and PG from IM proteoliposomes and PE, PG and POPC from OM proteoliposomes to apo-MlaC. Purified MlaC contains endogenous PL, which was removed by washing in detergent before used for transport reaction. All TLC experiments were done in the present of apo-MlaC. MlaA(∆Asn41-Phe42)/OmpF mutant was tested in the transport assay by TLC, showing inhibition to the retrograde PL transport. (H) SDS-PAGE analysis of proteins involved in the transported system of Fig. f. (I and J) FRET scan of PL transport assay using MlaA/OmpF wildtype or mutant MlaA(∆Asn41-Phe42)/OmpF in anterograde direction (I) or retrograde direction (J). The mutant abolished retrograde transport but not anterograde transport. (K) FRET scan of purified MlaC after PL transport reaction containing IM and OM proteoliposomes and apo-MlaC, measuring the fluorescent NB signal of the captured PE from anterograde or retrograde. (L and M) FRET scan of PL transport assay containing IM and OM proteoliposomes and MlaC in the presence or absence of ATP, $MgCl_2$ or AMP-PNP for anterograde (IM proteoliposomes contain fluorescent PE) (L) or retrograde (OM proteoliposomes contain fluorescent PE) (M) directions. FRET signals for both directions show to be ATP and $MgCl_2$ dependent. (N) FRET scan of PL transport assay containing IM proteoliposomes and apo-MlaC only in the presence or absence of ATP and $MgCl_2$. The result shows ATP independent anterograde in the presence of apo-MlaC, suggesting that ATP is required for releasing the bound PL in the complex. (O) FRET scan of purified OM proteoliposomes after PL transport reaction containing IM and OM with or without apo-MlaC. Result shows increased NB (530nm) and Rhod (590nm) signals after transport. (P) SDS-PAGE analysis of proteins in the purified OM proteoliposomes showing MlaA/OmpF and large amount of associated MlaC. Data are representative results from n$^{3}$3 experiments. Data in B-D, K represents mean ± s.d. (n$^{3}$3 experiments).

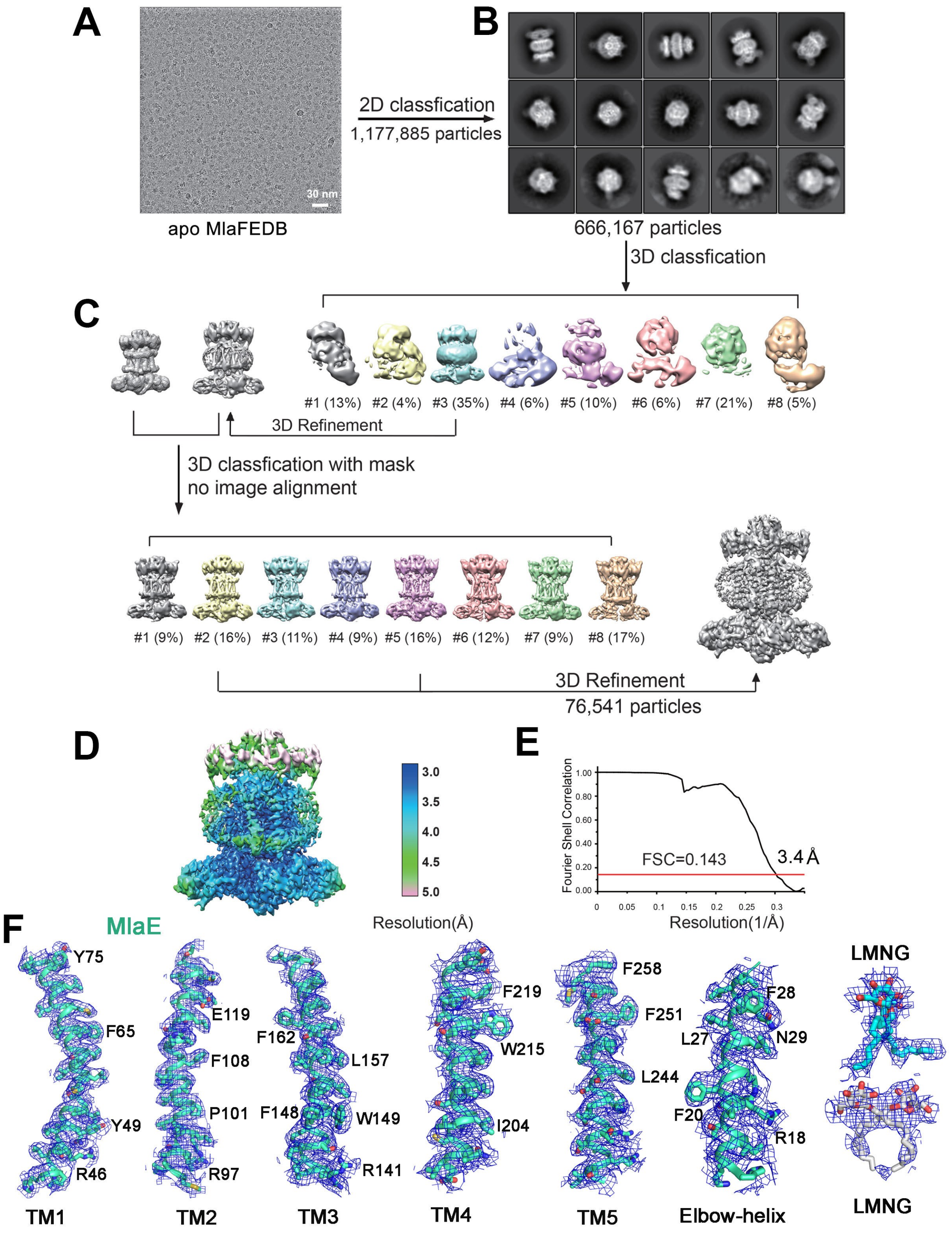


**Figure S2. Flowchart for cryo-EM single-particle data processing of native MlaFEDB complex.** (A) A micrograph of the single particles after drift correction and dose-weighting. (B) 2D classifications. (C) 3D classification, selections and 3D refinement. (D) The overall EM maps of apo MlaFEDB are colour coded to indicate the range of resolutions. (E) Gold-standard FSC curves of the final EM maps. (F) Residues with detailed side chains from TM1-TM5, and elbow helix of MlaE, and two LMNGs are shown in the cryo-EM map of apo MlaFEDB.

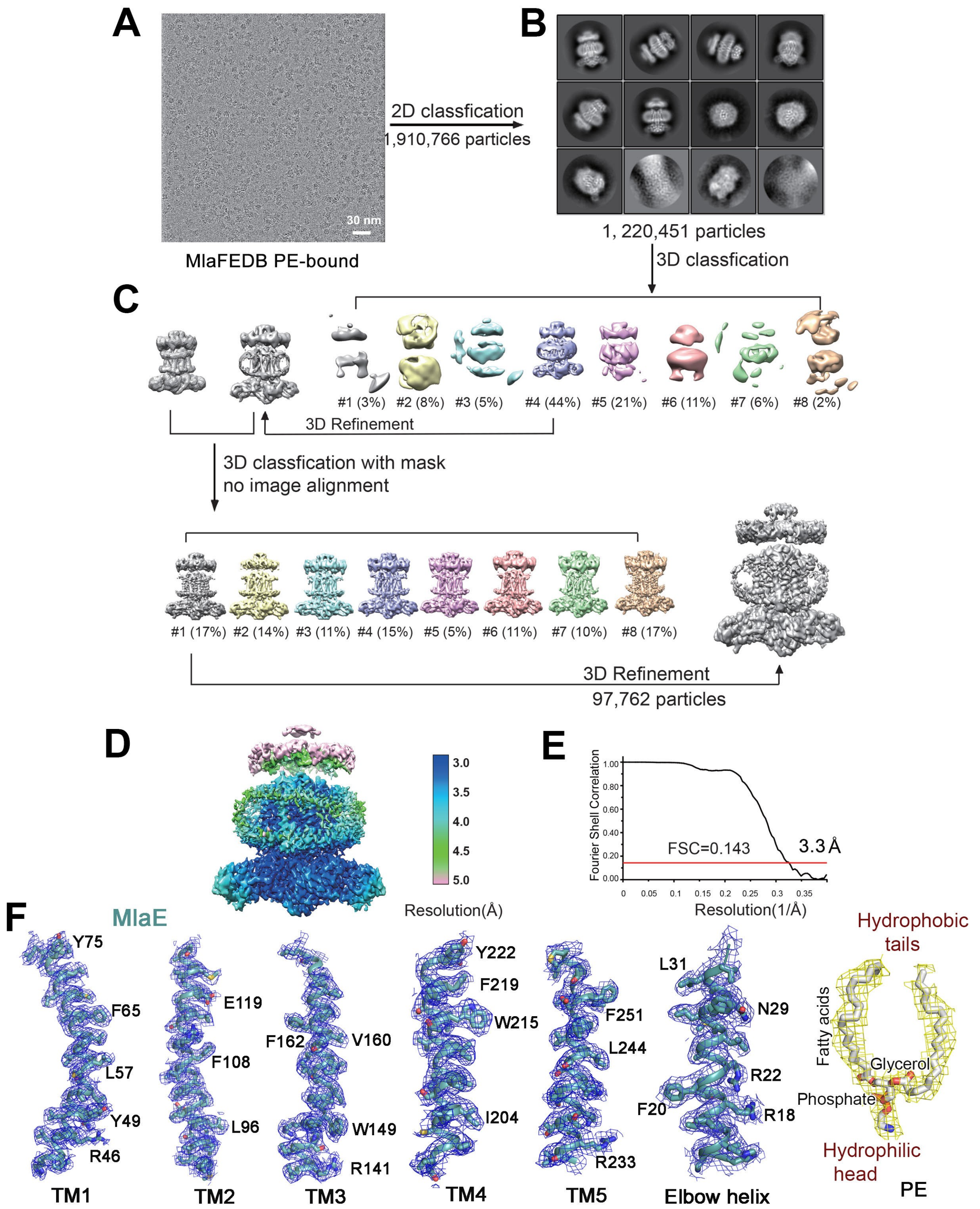


**Figure S3. Flowchart for cryo-EM single-particle data processing of PE bound MlaFEDB.** (A) A micrograph of the single particles after drift correction and dose-weighting. (B) 2D classifications. (C) 3D classification, selections and 3D refinement. (D) The overall EM map of the PL bound MlaFEDB is colour coded to indicate the range of resolutions. (E) The Gold-standard FSC curve of the final EM map. (F) Residues with detailed side chains of TM1-TM5 and elbow helix of MlaE, and a PE are shown in the cryo-EM map densities of PL-bound MlaFEDB complex.

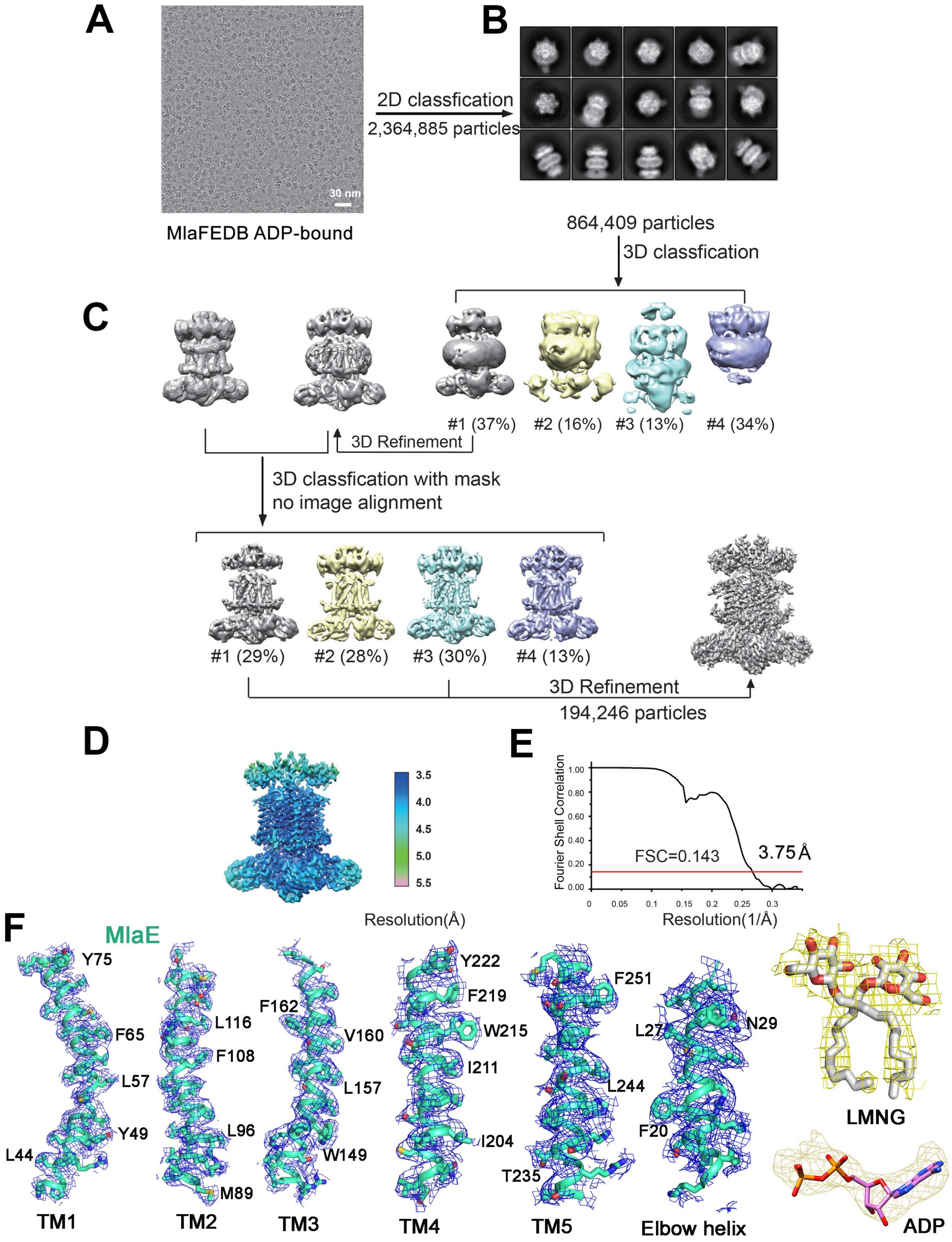


**Figure S4. Flowchart for cryo-EM single-particle data processing of ADP bound MlaFEDB.** (A) A micrograph of the single particles after drift correction and dose-weighting. (B) 2D classifications. (C) 3D classification, selections and 3D refinement. (D) The overall EM maps of ADP bound MlaFEDB are colour coded to indicate the range of resolutions. (E) The Gold-standard FSC curves of the final EM maps ofADP bound MlaFEDB. (F) Residue with detailed side chains of TM1-TM5 and the elbow helix of MlaE are shown in the cryo-EM map densities for the ADP-bound structure. LMNG and ADP are fitted for the extra density in the cavity and NBD.

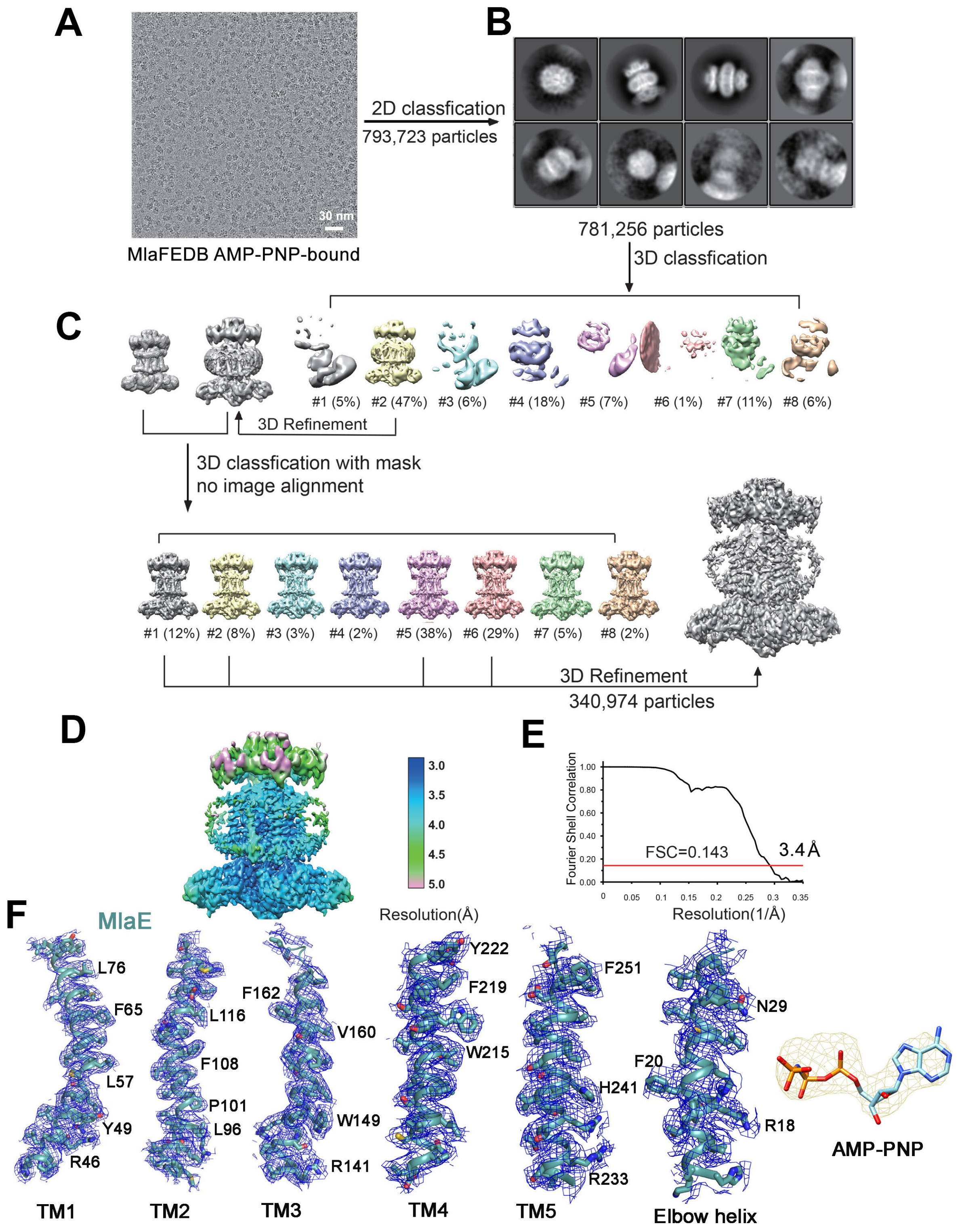


**Figure S5. Flowchart for cryo-EM single-particle data processing of AMP-PNP bound MlaFEDB.** (A) A micrograph of the single particles after drift correction and dose-weighting. (B) 2D classifications. (C) 3D classification, selections and 3D refinement. (D) The overall EM maps of AMP-PNP bound MlaFEDB are colour coded to indicate the range of resolutions. (E) The gold-standard FSC curves of the final EM maps. (F) Residue of TM1-TM5 and the elbow helix of MlaE are fitted in the map densities of AMP-PNP structure. AMP-PNP is fitted for the extra density in the NBD.

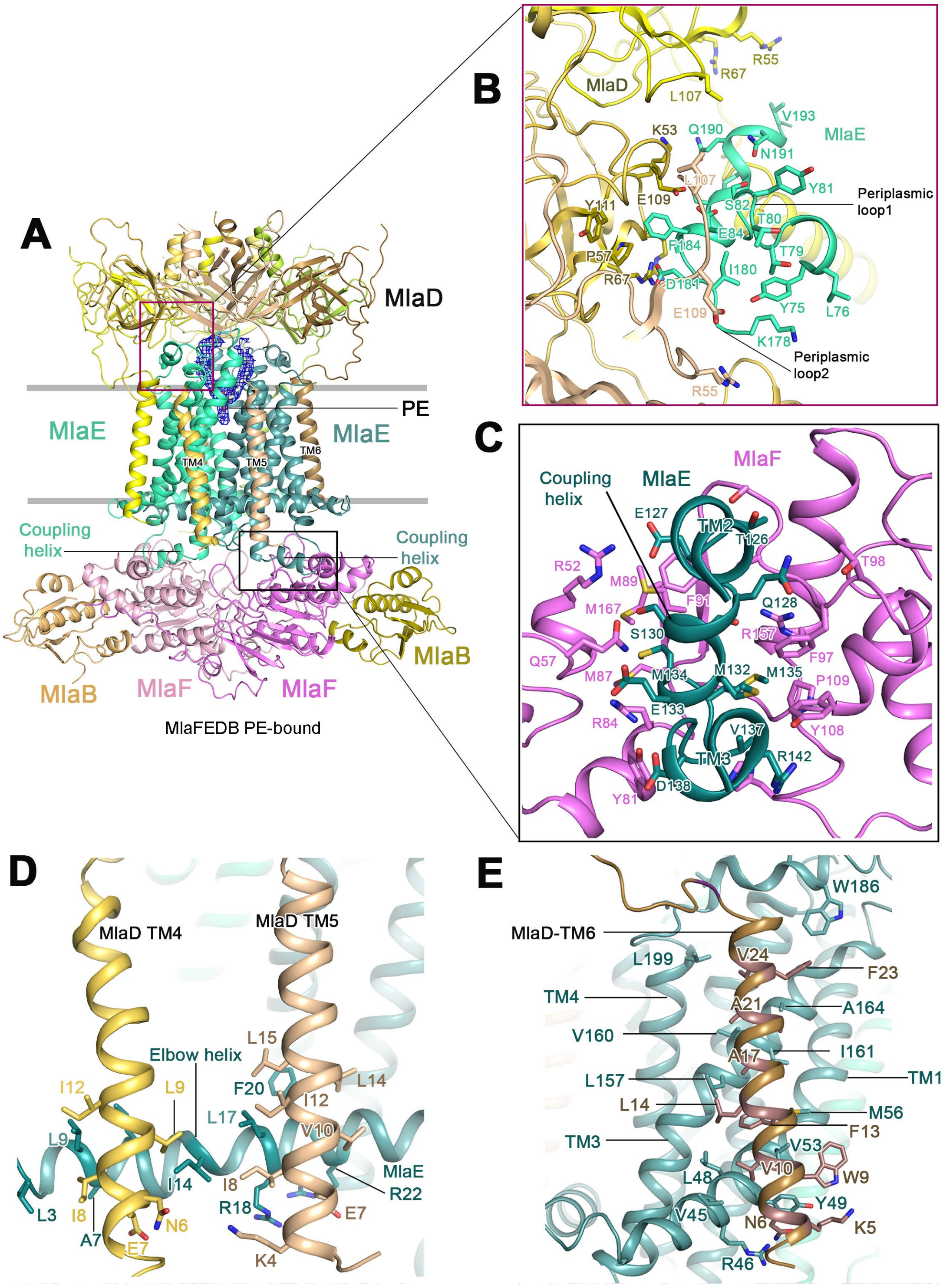


**Figure S6. Interactions of MlaE to the peripasmic MlaD and the cytoplasmic MlaF**. (A) Cartoon representation of PE-bound MlaFEDB complex. The colour scheme is the same as the Figure 2B. (B) MlaE interacts with the periplasmic domain of MlaD through the periplasmic loop 1 and 2. The interacting residues are labelled. (C) The coupling helix residues of one MlaE unit interact with residues located at the groove of one MlaF unit. (D) Two TM segments of MlaD interacting with the elbow helix of MlaE. (E) One TM segment of MlaD interacting with the TM1 and TM3 of MlaE.

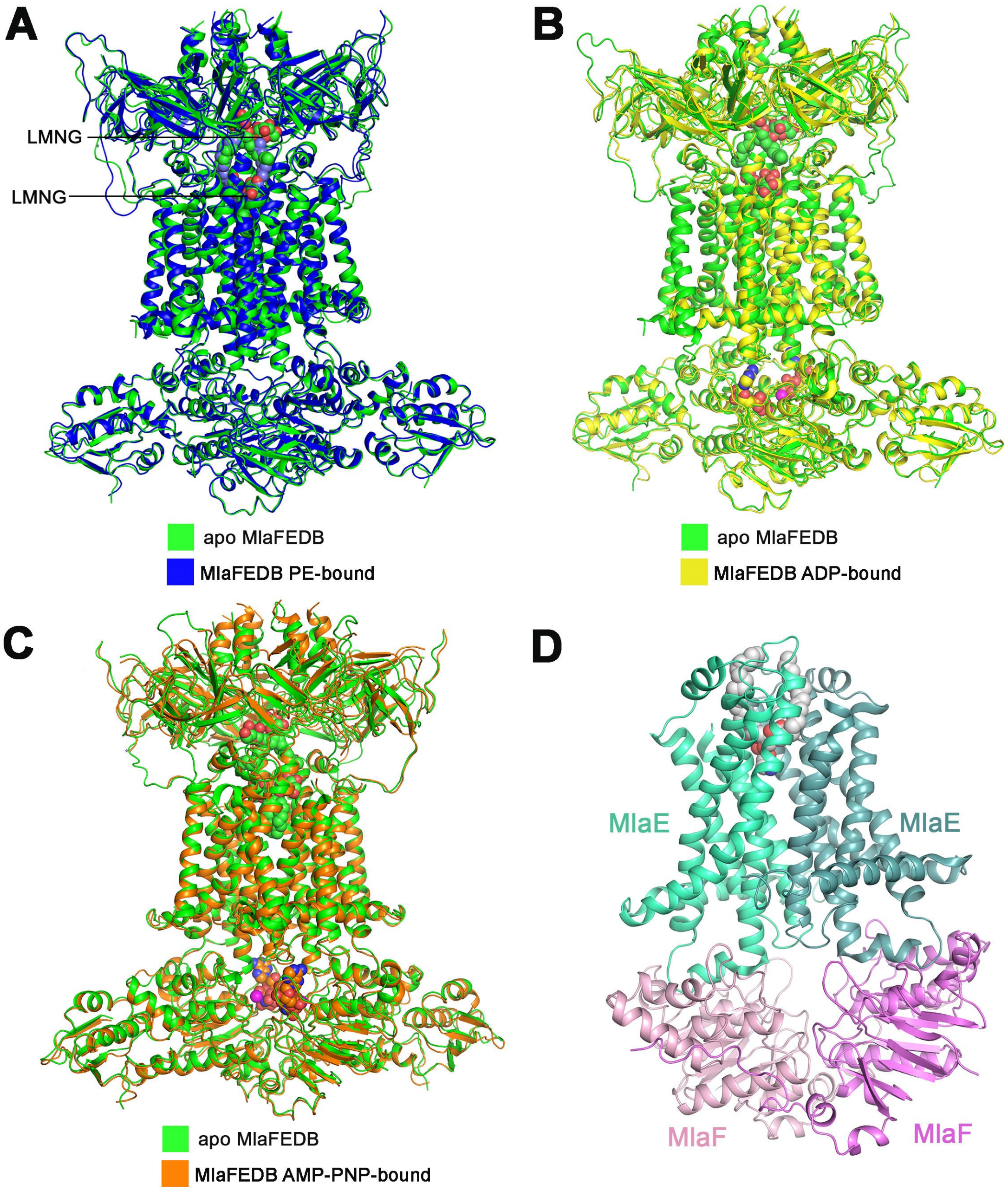


**Figure S7. Comparison of PE, ADP, AMP-PNP bound MlaFEDB structures to the apo structure of MlaFEDB**. The apo MlaFEDB is shown in green. The PL, ADP, AMP-PNP and LMNG are shown in sphere. (A) PE bound MlaFEDB complex is superimposed to the apo MlaFEDB complex. PE-bound MlaFEDB complex is coloured in blue. (B) ADP-bound MlaFEDB structure is superimposed to the apo MlaFEDB. ADP-bound MlaFEDB is coloured in yellow. (C) AMP-PNP bound MlaFEDB structure is superimposed to the apo MlaFEDB. AMP-PNP bound MlaFEDB is in orange. (D) Cartoon representation of MlaFE structure of the complex.

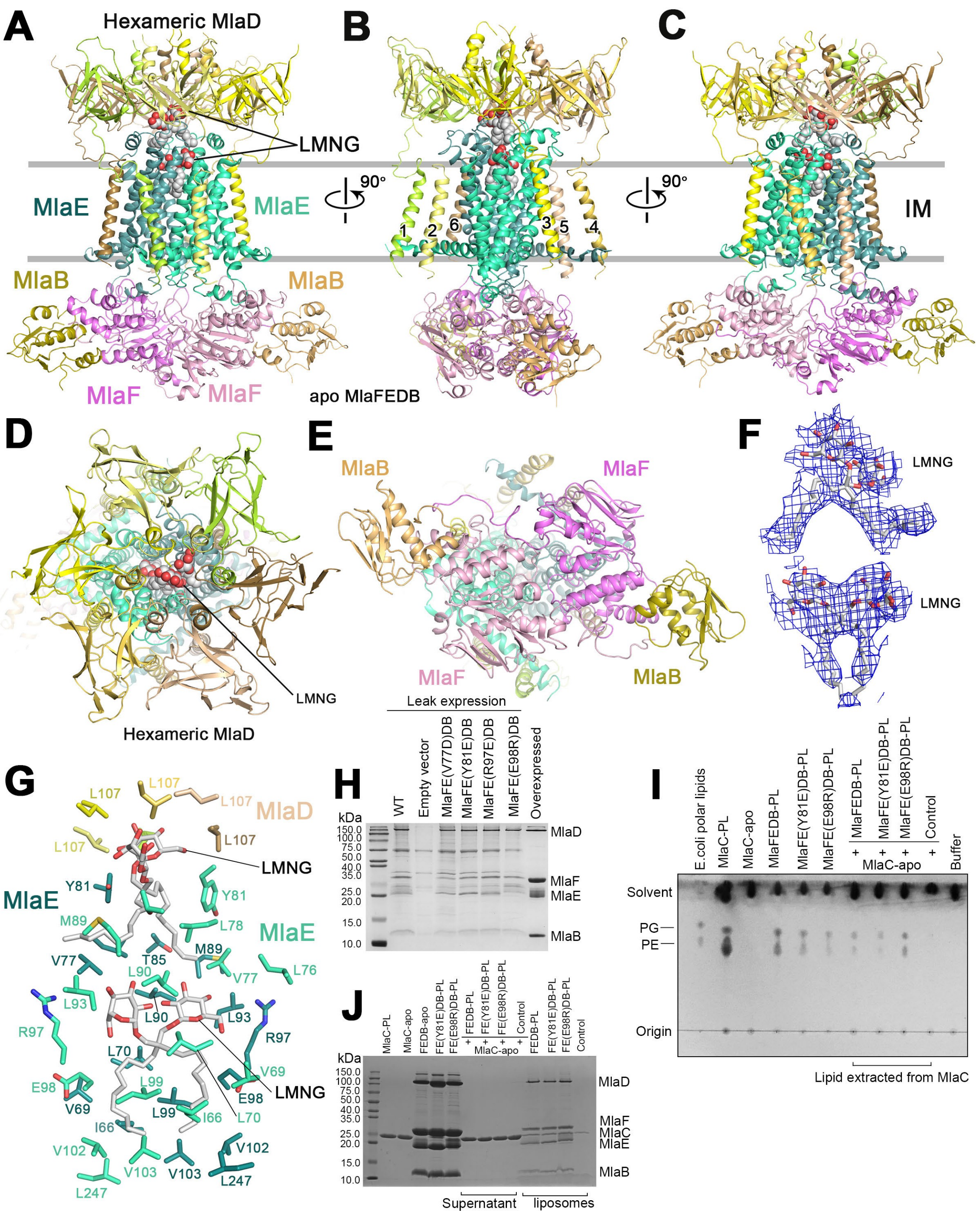


**Figure S8. MlaE ligand binding residues in the cavity**. (A) Cartoon representation of apo MlaFEDB. LMNG molecules are shown in sphere with carbon in grey and oxygen in red. (B) 90° rotation along the y-axis relative to the left panel. (C) 90° rotation of Fig. b along the y-axis. (D) Periplasmic view of apo MlaFEDB, showing hexameric MlaD. The LMNG molecule is seen through the MlaD channel. (E) Cytoplasmic view of native MlaFEDB. (F) Cryo-EM densities of LMNG molecules. (G) LMNG molecules are interacted with residues of MlaE and MlaD. MlaD residue L107, MlaE residues Y81, L78, V77, M89 interact the top LMNG molecule, while MlaE residues L90, R97, Q73, S94, L70, I66, V102, L247 and V103 interact with the bottom LMNG molecule.

(H) Leak expression of pTRC99a_MlaFEDCB in *E. coli* MlaE null strain, with or without MlaE mutations V77D, Y81E, E98R. (I) PL transport assay by TLC showing transported PL to apo-MlaC using wildtype MlaFEDB and mutant MlaFE(Y81E)DCB or MlaFE(E98R)DCB constituted IM proteoliposomes in the absence of ATP. Mutation of MlaE(E98R) affects the amount of endogenous PL binding in MlaFEDB. The results show reduced anterograde transport by the mutant MlaFE(Y81E)DCB and increased anterograde transport by the mutant MlaFE(E98R)DCB comparing with the wildtype MlaFEDB. (J) SDS-PAGE analysis of proteins involved in the PL transport assay of Figure I.

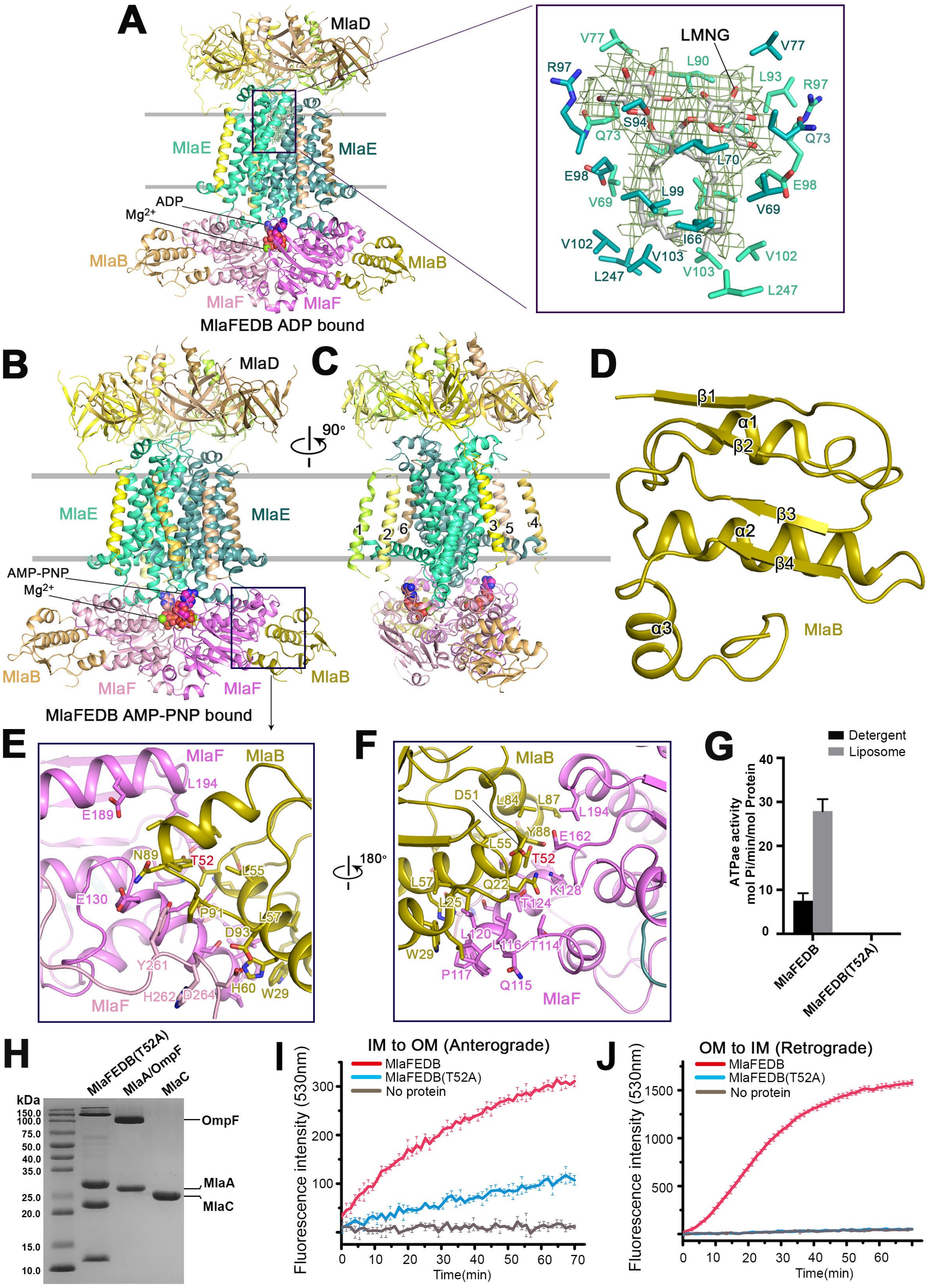


**Figure S9. Interactions of MlaB to MlaF using nucleotide bound structures of MlaFEDB**. (A) Cartoon representation of structure of the ADP-bound MlaFEDB complex, and the closed-view of the LMNG binding residues. The LMNG molecules is shown in stick and coloured in grey. Cryo-EM density of LMNG is on the right panel. The LMNG molecule is shown in stick and only two sugars are visible. (B) Cartoon representation of structure of the AMP-PNP bound MlaFEDB complex. The colour scheme is same as Figure A. (C) 90° rotation along the y-axis relative to the left panel. (D) Structure of MlaB showing the STAS domain, consisting of three *a*-helices and four *b*-strands. (E and F) The interactions between one MlaB and one MlaF involving the residues MlaB T52 (E) and 90° rotation along the y-axis relative to left panel (F). (G) Relative ATPase activity of MlaFEDB(T52A) in detergent or liposomes. (H) SDS-PAGE analysis of proteins of wildtype and mutant MlaFEDB(T52A). (I and J) FRET scan of PL transport using mutant MlaFEDB(T52A) or wildtype MlaFEDB IM proteoliposomes in the presence of ATP and $MgCl_2$ for anterograde (I) or retrograde (J) direction. The result show that the mutation only abolished retrograde signal but not anterograde signal.

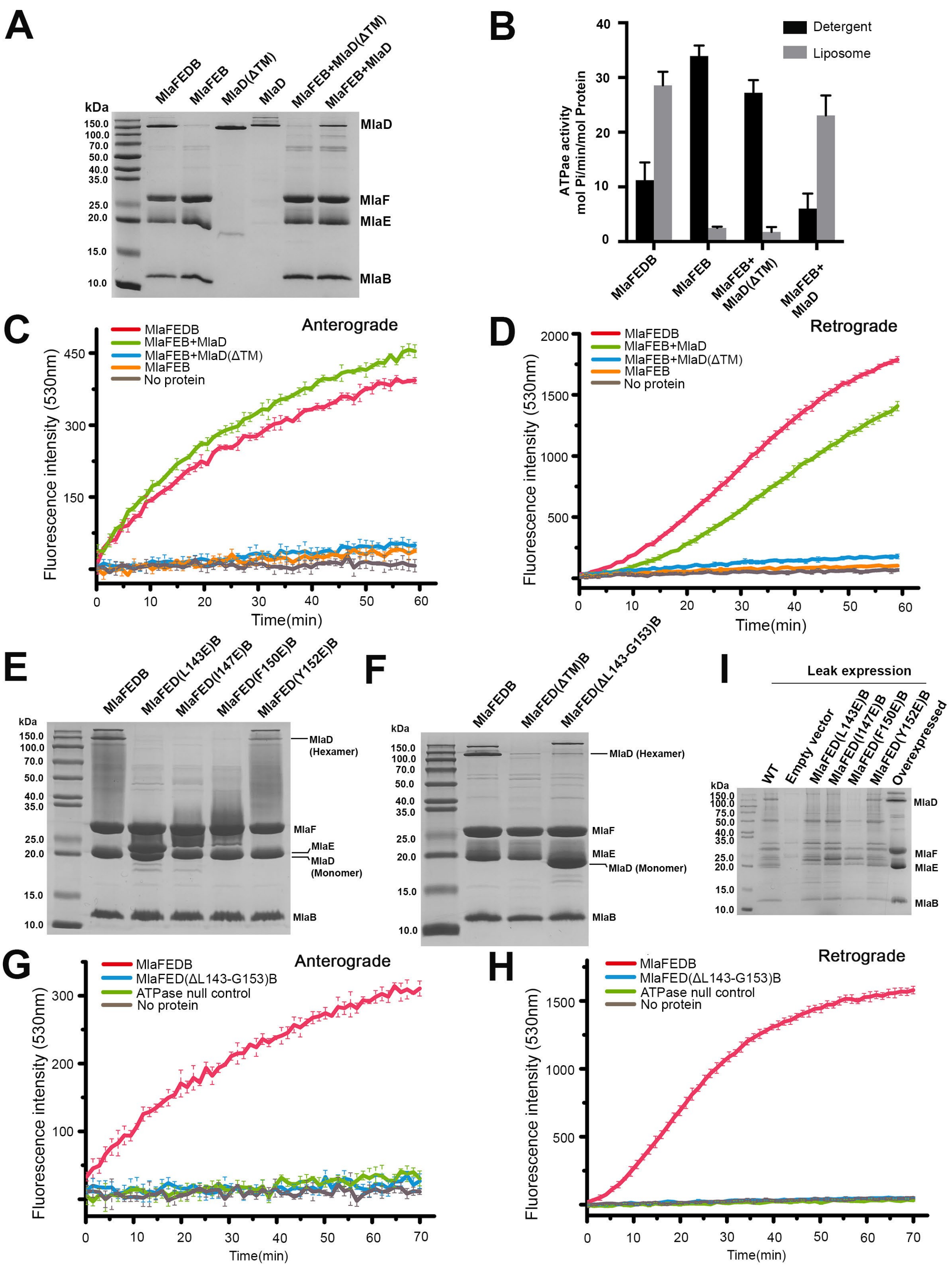


**Figure S10. Functional residues characterization of MlaD**. (A) Coomassie brilliant blue staining of purified Mla proteins. Purified MlaFEB was incubated with TM truncated MlaD(ΔTM) or full length MlaD to form complex *in vitro*, which were purified using affinity columns and analysed on SDS-PAGE along with wild-type MlaFEDB and MlaFEB. (B) The relative ATPase activities of wild-type, MlaFEB and *in vitro* formed MlaFEDB were measured in detergent and liposomes. Soluble MlaD(ΔTM) was added into MlaFEB constructed systems and its effect to ATPase activity was also analysed. (C and D) FRET scan of PL transport assay using wildtype MlaFEDB or MlaFEB complex incubated with truncated MlaD(ΔTM) or full length MlaD for anterograde direction (C) or retrograde direction (D). Soluble MlaD(ΔTM) was added into MlaFEB constructed systems and its effect to transport activity was also analysed. (E) SDS-PAGE analysis of purified MlaFEDB complexes with MlaD mutation I143E, I147, F150E or Y152E. The mutants MlaD I143E, I147E and F150E lost the SDS-resistance hexameric form. (F) SDS-PAGE analysis of truncated mutants including MlaD TM segments deleted MlaFED(ΔTM)B, MlaD MCE channel and deleted MlaFED(ΔL143-G153)B. (G and H) FRET scan of PL transport assay using wildtype MlaFEDB or MlaFEDB mutants MlaFED(ΔL143-G153)B for anterograde (G) or retrograde (H) direction. (I) Leak expression of pTRC99a_MlaFEDCB containing MlaD mutation L143E, I147E, F150E or Y152E. Data are representative results from n ≥3 experiments. Data in B represents mean ± s.d. (n ≥3);

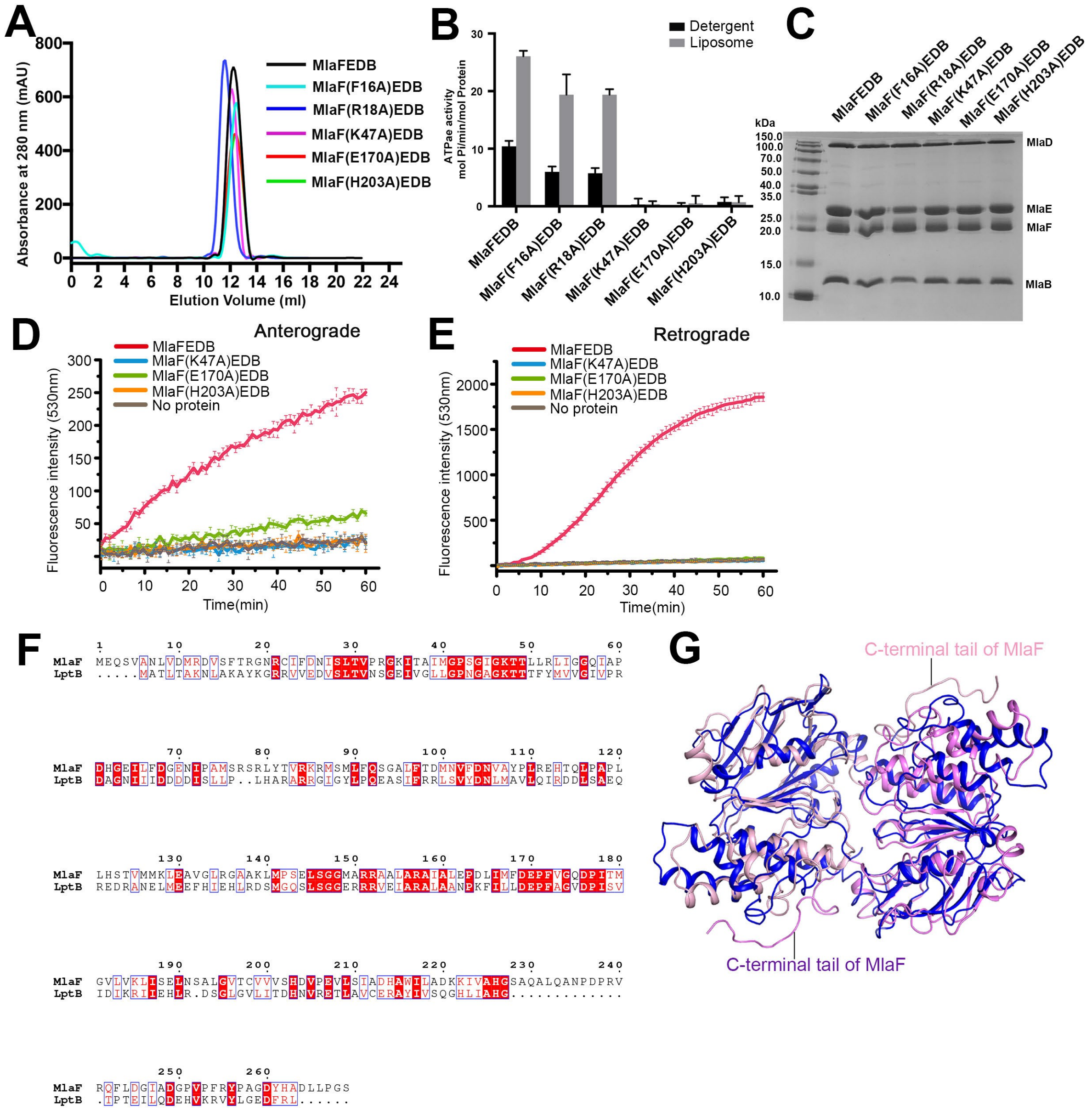


**Figure S11. Effect of nucleotide binding residues of MlaF mutation on ATPase activity and *in vitro* PL transport**. (A) The size-exclusion chromatogram of purified MlaFEDB complexes with MlaF single catalytic mutants of F16A, R18A, K47A, E170A and H203A. The mutants were eluted almost at the same time as the wild-type MlaFEDB, indicating that the mutants have the similar structure as the wild-type. (B) The relative ATPase activities of MlaFEDB wild-type and mutants in detergent and liposomes. (C) SDS-PAGE analysis of purified wildtype and the mutants. (D and E) FRET scan of PL transport using MlaFEDB mutant IM proteoliposomes containing MlaF K47A, E170A or H203A, wildtype OM proteoliposomes and MlaC for anterograde (D) or retrograde (E). The catalytic and ATP binding residue mutants abolished retrograde transport while the catalytic mutant MlaF(E170A)EDB does not abolish anterograde transport of PLs. (F) Amino acid sequence alignment of MlaF and LptB. MlaF and LptB have 24.54% amino acid identity. The C-terminal tail of MlaF is longer than that of LptB. (G) Dimeric MlaF is superimposed into the dimeric LptB. MlaF has a long C-terminal tail that interacts with the other MlaF molecule, but LptB does not. Data are representative results from n$^{3}$3 experiments. Data in B represents mean ± s.d. (n$^{3}$3 experiments).

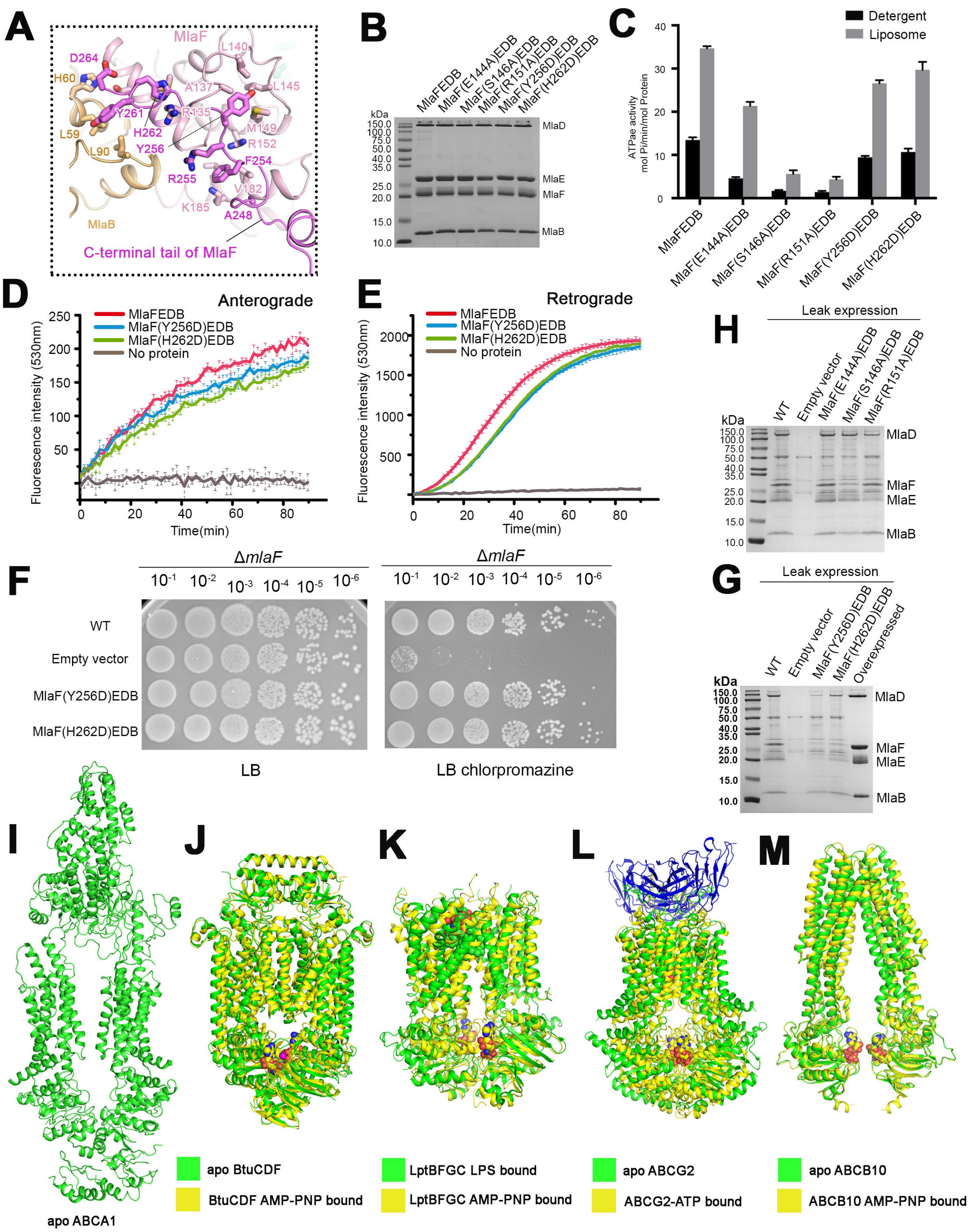

**Figure S12. Functional interacting residues between MlaB and MlaF**. (A) MlaF C-terminal tail showing interacting residues with the opposite MlaF and MlaB. (B) SDS-PAGE analysis of purified MlaFEDB wildtype or mutant containing single mutation of the signature motif residues mutants E144A, S146S, R151A of MlaF, and C-terminal residues Y256D and H262D of MlaF. (C) The relative ATPase activity of MlaFEDB mutants in detergent and in liposomes. Signature mutants significantly reduce the ATPase activities, but Y256D and H262D mutants do not change the ATPase activity.

(D and E) FRET scan of PL transport assay using wildtype MlaFEDB or mutant MlaF(Y256D) or MlaF(H262D) IM proteoliposomes, wildtype OM proteoliposome and MlaC for anterograde for anterograde (D) or retrograde (E) direction. h, Leak expression of pTRC99a_MlaFEDCB containing the mutations. (F) Cellular sensitivity to chlorpromazine by the mutant MlaF(Y256D)EDB or MlaF(H262D)EDB, showing no cellular effect. (G) Leak expression of pTRC99a_MlaF(Y256D)EDB or MlaF(H262D)EDB. (H) Leak expression of pTRC99a_MlaFEDCB containing MlaF mutation E144A, S146A, and R151A. Data are representative results from n ≥3 experiments. Data in C represents mean ± s.d. (n ≥3). (I-M) Structures of reported ABC transporters in apo and AMP-PNP bound states, (I) Human ABCA1 lipid exporter, apo ABCA1 (PDB code: 5XJY). The ABCA1 has limited structural similarity to MlaFEDB (Dali server search with a Z score of 10). (J) *E. coli* vitamin B12 importer, apo BtuCDF (PDB code: 2QI9), BtuCDF in complex with AMP-PNP (PDB code: 4FI3). BtuCDF binding AMP-PNP causes conformational changes. (K) *E. coli* lipopolysaccharide transporter, apo LptBFGC (PDB code: 6S8N), LptBFGC in complex with AMP-PNP (PDB code: 6S8G). LptB has some structural similarity to MlaF. LptBFGC binding AMP-PNP causes significant conformational changes. (L) Human ABCG2 multidrug transporter, apo (PDB code: 6MIJ), ABCG2 E211Q mutant in complex with ATP (PDB code: 6HZM). ATP binding ABCG2 causes conformational changes. (M) Human mitochondrial ABC transporter ABCB10, apo (PDB code: 3ZDQ), in complex with AMP-PNP (PDB code: 4AYW). The AMP-PNP binding does not cause significant conformational changes.

**Table S1** | Statistics of Cryo-EM data collection and structure refinement

| | MlaFEDB native | MlaFEDB PE-bound | MlaFEDB AMP-PNP-bound | MlaFEDB ADP-bound |
|---|---|---|---|---|
| **Data Collection** | | | | |
| EM equipment | Titan Krios (Thermo Fisher) | | | |
| Magnification | 49310 | | | |
| Voltage (kV) | 300 | | | |
| Detector | Gatan K2 Summit | | | |
| Pixel size (Å) | 1.014 | | | |
| Electron dose (e-/Å2) | 56 | | | |
| Defocus range (μm) | -1.0 ~ -2.8 | | | |
| **Reconstruction** | | | | |
| Software | RELION 3.0 | | | |
| Number of used particles | 76,541 | 97,762 | 340,974 | 194,246 |
| Symmetry | C2 | | | |
| Final Resolution (Å) | 3.4 | 3.3 | 3.4 | 3.75 |
| Map sharpening B-factor (Å2) | -85 | -77 | -172 | -182 |
| **Refinement** | | | | |
| Software | Phenix | Phenix | Phenix | Phenix |
| Model composition | | | | |
| Protein residues | 2129 | 2102 | 2066 | 2016 |
| Side chains assigned | 2129 | 2102 | 2066 | 2016 |
| AMP-PNP | 0 | 0 | 2 | 0 |
| ADP | 0 | 0 | 0 | 2 |
| Detergents | 2 | 0 | 0 | 1 |
| PL | 0 | 1 | 0 | 0 |
| $Mg^{2+}$ | 0 | 0 | 2 | 2 |
| **R.m.s deviations** | | | | |
| Bonds length (Å) | 0.011 | 0.007 | 0.007 | 0.007 |
| Bonds Angle (°) | 1.528 | 1.105 | 1.086 | 1.212 |
| **Ramachandran plot statistics** | | | | |
| Preferred (%) | 84.06 | 87.67 | 86.04 | 84.50 |
| Allowed (%) | 15.23 | 12.04 | 13.46 | 14.29 |
| Outlier (%) | 0.71 | 0.29 | 0.50 | 1.21 |
